# PriSeT: Efficient *De Novo* Primer Discovery

**DOI:** 10.1101/2020.04.06.027961

**Authors:** Marie Hoffmann, Michael T. Monaghan, Knut Reinert

**Affiliations:** Institute of Computer Science, Freie Universität Berlin, Takustr. 9, 14195 Berlin, Germany; Institute for Biology, Freie Universität Berlin, Königin-Luise-Str. 1-3, 14195 Berlin, Germany; Leibniz Institute of Freshwater Ecology and Inland Fisheries (IGB), Müggelseedamm 301, 12587 Berlin, Germany

## Abstract

**Motivation:** DNA metabarcoding is a commonly applied technique used to infer the species composition of environmental samples. These samples can comprise hundreds of organisms that can be closely or very distantly related in the taxonomic tree of life. DNA metabarcoding combines polymerase chain reaction (PCR) and next-generation sequencing (NGS), whereby a short, homologous sequence of DNA is amplified and sequenced from all members of the community. Sequences are then taxonomically identified based on their match to a reference database. Ideally, each species of interest would have a unique DNA *barcode*. This short, variable sequence needs to be flanked by relatively conserved regions that can be used as primer binding sites. Appropriate PCR primer pairs would match to a broad evolutionary range of taxa, such that we only need a few to achieve high taxonomic coverage. At the same time however, the DNA barcodes between primer pairs should be different to allow us to distinguish between species to improve resolution. This poses an interesting optimization problem. More specifically: Given a set of references ℛ = {*R*_1_, *R*_2_, …, *R*_*m*_}, the problem is to find a primer set *P* balancing both: high taxonomic coverage and high resolution. This goal can be captured by filtering for frequent primers and ranking by coverage or variation, i.e. the number of unique barcodes. Here we present the software PriSeT, an offline primer discovery tool that is capable of processing large libraries and is robust against mislabeled or low quality references. It tackles the computationally expensive steps with linear runtime filters and efficient encodings.

**Results:** We first evaluated PriSeT on references (mostly 18S rRNA genes) from 19 clades covering eukaryotic organisms that are typical for freshwater plankton samples. PriSeT recovered several published primer sets as well as additional, more chemically suitable primer sets. For these new sets, we compared frequency, taxon coverage, and amplicon variation with published primer sets. For 11 clades we found *de novo* primer pairs that cover more taxa than the published ones, and for six clades *de novo* primers resulted in greater sequence (i.e., DNA barcode) variation. We also applied PriSeT to 19 SARS-CoV-2 genomes and computed 114 new primer pairs with the additional constraint that the sequences have no co-occurrences in other taxa. These primer sets would be suitable for empirical testing.

**Availability:** https://github.com/mariehoffmann/PriSeT

## 1. Motivation

In metagenomics, the DNA of hundreds to thousands of species from a single environmental sample is processed in parallel and one goal is to identify all species from a sample. Samples contain many species in what can be very heterogeneous communities.

Their identification is usually done via DNA *metabarcoding* which combines polymerase chain reaction (PCR), next-generation sequencing (NGS) and identification via DNA barcodes, i.e. short sequences that are matched against a reference database. In contrast, single DNA probing and assembly would be more precise, but are for many reasons not feasible, e.g., the majority of microorganisms cannot be clonally cultured – a necessity for assembling reference genomes (Rappé & Giovannoni, 2003).

Although the decreasing costs of NGS makes DNA metabarcoding progressively feasible, identification of most microeukaryotes, such as plankton, continues to be done via morphology. This still outperforms molecular methods in some aspects, like abundance estimation, differentiation of life stages, or observation of teratological forms. But metabarcoding of environmental DNA (eDNA) is a far more sensitive method of species detection compared to traditional methods (Smart et al., 2015) provided that a suitable marker is chosen.

In the search for a marker, we know that evolutionary pressure is not consistent within genomes. There are regions that vary even within species, or on the other extreme, are conserved for whole clades. For an effective delineation of taxa using DNA metabarcodes, we are interested in regions that are similar or identical to individuals of the same species, but distinctive from other species. Such regions are ideal barcodes, but they need to be surrounded by conserved regions that contain suitable binding sites for primers. Primers are short oligomers, ideally 16 to 25 bases long, which determine the copy starting point of the template. We need flanking conservation, because we batch-process all extracted DNA at once and can add only one primer pair per PCR run^1^.

The most frequently applied type of NGS in DNA metabarcoding is the paired-end approach as it allows for longer sequences and error correction for the low quality ends of reads. The two types of primers operate independently, one binds on the forward and the other on the reverse strand of the template. Both are elongated enzymatically to produce a complementary copy. In each PCR cycle the number of amplicons is nearly duplicated.

After about 25-35 cycles the reaction is stopped and the amplified DNA amplicons provide a signal for the sequencer machine that is separable from the noise of the original DNA material. After digitization of the reads, an analysis pipeline combines trimming, quality filtering, denoising, and read merging with the aforementioned error correction. Identical or similar sequences are then clustered into *operational taxonomic units* (OTUs). The underlying hypothesis is that each OTU corresponds to a taxon in the tree of life. An alternative approach is that each sequence variant is analysed rather than being clustered into OTUs. The art herein is to discriminate sequencing errors from variants. A noteworthy denoising algorithm is the Divisive Amplicon Denoising Algorithm 2 (DADA 2) (Callahan et al., 2016).

As described earlier, not all barcodes are specific to a single species and our metabarcoding pipeline will most likely produce OTUs uniting barcodes from taxonomically distinct lineages, even with similarity thresholds set high (i.e., > 97 %). Despite these natural imperfections, ecologists found that the small subunit RNA gene (SSU) is an excellent source for species-specific markers. Hence a large number of entries in the NCBI-GenBank nucleotide database originate from ribosomal subunits (Benson et al., 2012). Schmidt et al. (2014) demonstrated that OTUs from 16S/18S rRNA reflect the underlying ecological diversity consistently across habitats. In other words, even if OTUs cannot be resolved, their counts correlate to the sample’s diversity and are roughly independent from the species composition.

Finding the optimal set of primers in terms of coverage and resolution takes many costly iterations of trial and error (Elbrecht et al., 2019). Scientists encounter problems like missing ground truth and sparsely populated reference databases. Often they would start with primer sequences published in previous studies on similar data sets and then compare OTU diversity between different primer sets.

More formally, primer candidates can be described as *k*-mers – DNA oligomers of length *k* – satisfying a set of sequence constraints *C*_*s*_ which are relevant for the success of the PCR and which must hold independent of their matching. *C*_*s*_ comprises range limits for melting temperature, CG content or low probabilities for self-annealing. When combining two *k*-mers to produce a primer pair, both have to fulfill a second set of constraints *C*_*p*_ arising from their strand orientation and fitting: avoidance of CG clamps or low probability of cross-annealing.

A reference sequence of length *N* contains 𝒪(*KN* ^2^) many *k*-mer pairs which need to undergo filtering of *C*_*s*_ and *C*_*p*_ where *K* is the *k*-mer length range. We use the *k*-mer frequency computation on the FM index of the library to quickly identify frequent *k*-mers – above a threshold *F* – which then undergo two processing steps: filtering and combination into pairs.

## 2. Motivation

In metagenomics, the DNA of hundreds to thousands of species from a single environmental sample is processed in parallel and one goal is to identify all species from a sample. Samples contain many species in what can be very heterogeneous communities.

Their identification is usually done via DNA *metabarcoding* which combines polymerase chain reaction (PCR), next-generation sequencing (NGS) and identification via DNA barcodes, i.e. short amplicons that are matched against a reference database. In contrast, single DNA probing and assembly would be more precise, but are for many reasons not feasible, e.g., the majority of microorganisms cannot be clonally cultured – a necessity for assembling reference genomes (Rappé & Giovannoni, 2003).

Although the decreasing costs of NGS makes DNA metabarcoding progressively feasible, identification of most microeukaryotes, such as plankton, continues to be done via morphology. This still outperforms molecular methods in some aspects, like abundance estimation, differentiation of life stages, or observation of teratological forms. But metabarcoding of environmental DNA (eDNA) is a far more sensitive method of species detection compared to traditional methods (Smart et al., 2015) provided that a suitable marker is chosen.

In the search for a marker, we know that evolutionary pressure is not consistent within genomes. There are regions that vary even within species, or on the other extreme, are conserved for whole clades. For an effective delineation of taxa using DNA metabarcodes, we are interested in regions that are similar or identical to individuals of the same species, but distinctive from other species. Such regions are ideal barcodes, but they need to be surrounded by conserved regions that contain suitable binding sites for primers. Primers are short oligomers, ideally 16 to 25 bases long, which determine the copy starting point of the template. We need flanking conservation, because we batch-process all extracted DNA at once and can add only one primer pair per PCR run^2^.

The most frequently applied type of NGS in DNA metabarcoding is the paired-end approach as it allows for longer sequences and error correction for the low quality ends of reads. The two types of primers operate independently, one binds on the forward and the other on the reverse strand of the template. Both are elongated enzymatically to produce a complementary copy. In each PCR cycle the number of amplicons is nearly duplicated.

After about 25-35 cycles the reaction is stopped and the amplified DNA amplicons provide a signal for the sequencer machine that is separable from the noise of the original DNA material. After digitization of the reads, an analysis pipeline combines trimming, quality filtering, denoising, and read merging with the aforementioned error correction. Identical or similar sequences are then clustered into *operational taxonomic units* (OTUs). The underlying hypothesis is that each OTU corresponds to a taxon in the tree of life. An alternative approach is that each sequence variant is analysed rather than being clustered into OTUs. The art herein is to discriminate sequencing errors from variants. A noteworthy denoising algorithm is the Divisive Amplicon Denoising Algorithm 2 (DADA 2) (Callahan et al., 2016).

As described earlier, not all barcodes are specific to a single species and our metabarcoding pipeline will most likely produce OTUs uniting barcodes from taxonomically distinct lineages, even with similarity thresholds set high (i.e., > 97 %). Despite these natural imperfections, ecologists found that the small subunit RNA gene (SSU) is an excellent source for species-specific markers. Hence a large number of entries in the NCBI-GenBank nucleotide database originate from ribosomal subunits (Benson et al., 2012). (Schmidt et al., 2014) demonstrated that OTUs from 16S/18S rRNA reflect the underlying ecological diversity consistently across habitats. In other words, even if OTUs cannot be resolved, their counts correlate to the sample’s diversity and are roughly independent from the species composition.

Finding the optimal set of primers in terms of coverage and resolution takes many costly iterations of trial and error (Elbrecht et al., 2019). Scientists encounter problems like missing ground truth and sparsely populated reference databases. Often they would start with primer sequences published in previous studies on similar data sets and then compare OTU diversity between different primer sets.

More formally, primer candidates can be described as *k*-mers – DNA oligomers of length *k* – satisfying a set of sequence constraints *C*_*s*_ which are relevant for the success of the PCR and which must hold independent of their matching. *C*_*s*_ comprises range limits for melting temperature, CG content or low probabilities for self-annealing. When combining two *k*-mers to produce a primer pair, both have to fulfill a second set of constraints *C*_*p*_ arising from their strand orientation and fitting: avoidance of CG clamps or low probability of cross-annealing.

A reference sequence of length *N* contains 𝒪(*KN* ^2^) many *k*-mer pairs which need to undergo filtering of *C*_*s*_ and *C*_*p*_ where *K* is the *k*-mer length range. We use the *k*-mer frequency computation on the FM index of the library to quickly identify frequent *k*-mers – above a threshold *F* – which then undergo two processing steps: filtering and combination into pairs.

### 2.1. Existing Approaches

There exists computer-assisted approaches in which a manageable subset of references of organisms that are expected to show up, is collected and serves as input to compute a *multiple sequence alignment* (MSA). Then a variation or entropy score is computed given the nucleotide distribution for each position of the alignment. Low entropy regions are then analyzed for serving as primer templates and regions with high inter-species entropy as barcodes (Hadziavdic et al., 2014). MSA approaches require the input data set to be small, phylogenetically close, and curated in order to produce meaningful alignments. Needless to say that in metagenomics this approach is only chosen *post experimentum* when analysing organism groups that turned out to be difficult to detect with PCR.

Online tools like Primer Blast/Primer3^3^ (Ye et al., 2012) take off the burden to locally set up a software system by providing primer searches as on online service. Users provide as input a GenBank accession or a FASTA file. NCBI’s Primer BLAST uses Primer3 to design primer sequences and then submits a BLAST search against a user-provided file. Surprisingly, Primer Blast does not process files with multiple references and is therefore inapplicable in a metabarcoding setup.

The Primer Search Tool^4^ by Tusnady et al. (2005) uses a dynamic programming approach inspired by Kaempke et al. (2001). When combining candidate sequences Kaempke et al. (2001) make use of the sequence overlap and re-use computations of the shared substring. The online tool allows the upload of files containing multiple references, but produces errors when header lines contain special characters or sequences contain ambiguous one-letter encodings. A file of 114 KB sequences took 40 seconds to be processed including network traffic^5^. It outputs primer pairs scored by chemical fitness, but does not optimize towards coverage or amplicon variation.

PRIMEX (Lexa & Valle, 2004) is intended to provide querying as an online service. The tool performs internally a lookup of *k*-mers derived from the query string in a prepared library and allows for mismatches. Unfortunately, PRIMEX is currently not available and it is not capable of discovering new primer sequences^6^.

One of the few standalone tools is FastPCR (Kalendar et al., 2017). It supports a variety of PCR protocols and chemical checks. It first computes a hash table with *k*-mers grouped together by overlap into so called *k*-tuples with *k* being 7, 9, or 12 nt long. The derivation of these *k*-tuples is not described. In a second step, FastPCR uses a sliding window approach to search for matches between *k*-tuples and reference sequences. Upon match, individual *k*-tuples are extended in both directions. FastPCR has no preference for frequent *k*-mers – it is designed to work on single genomes. The authors are not aware of a primer discovery tool suitable for metabarcoding experiments where the aim is a high taxonomic coverage and barcode variation and the input is a directory or source file composed of thousands of reference sequences. Before we explain in detail the data structures and algorithm to solve the problem of primer discovery, we give definitions of some concepts that are essential to make our approach time-efficient.

#### Definition *k*-mer

With *k*-mer we refer to a sub-sequence of length *k* as part of a biological text *T*. A sequence of length *n* contains *n* − *k* + 1 *k*-mers. For an arbitrary alphabet Σ, there exist |Σ |^*k*^ different *k*-mers, i.e. for DNA 4^*k*^ unique *k*-mers. Since DNA is inherently structured, we do not expect *k*-mers to be uniformly distributed.

#### Definition (*k, e*)-frequency

The (*k, e*)-frequency counts for each of the *n* − *k* + 1 *k*-mers its frequency within the text *T* with up to *e* errors. For example, the (4, 0)-frequency of *T* = AACGACGATGCAGTACGAT over Σ = {`A;C;G;T`} is^7^:

|  |  |  |  |  |  |  |  |  |  |  |  |  |  |  |  |  |
| --- | --- | --- | --- | --- | --- | --- | --- | --- | --- | --- | --- | --- | --- | --- | --- | --- |
| $i$ | 0 | 1 | 2 | 3 | 4 | 5 | 6 | 7 | 8 | 9 | 10 | 11 | 12 | 13 | 14 | 15 |
| $T[i]$ | A | A | C | G | A | C | G | A | T | G | C | A | G | T | A | C |
| $F_{4,0}[i]$ | 1 | 3 | 1 | 1 | 3 | 2 | 1 | 1 | 1 | 1 | 1 | 1 | 1 | 1 | 3 | 2 |

#### Definition FM-index

The FM-index is a substring index over a text. It is based on the Burrows-Wheeler transform and adds support data structures that facilitate navigation when searching for query strings. Its efficiency comes from the adjacent location of substrings having prefixes in common. A word of length *k* can be looked up in 𝒪(*k*) (Ferragina & Manzini, 2000).

#### Definition rank

Given a text index *i*, the **rank** of a symbol *σ* ∈ Σ, outputs the number of letters equal to *σ* until position *i*, including the text position *i* itself.

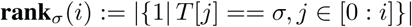

Example: **rank**_1_ of *T* = `100101111010001` over Σ = {0, 1}

|  |  |  |  |  |  |  |  |  |  |  |  |  |  |  |  |
| --- | --- | --- | --- | --- | --- | --- | --- | --- | --- | --- | --- | --- | --- | --- | --- |
| $i$ | 0 | 1 | 2 | 3 | 4 | 5 | 6 | 7 | 8 | 9 | 10 | 11 | 12 | 13 | 14 |
| $T[i]$ | 1 | 0 | 0 | 1 | 0 | 1 | 1 | 1 | 1 | 0 | 1 | 0 | 0 | 0 | 1 |
| **rank**<sub>1</sub>[$i$] | 1 | 1 | 1 | 2 | 2 | 3 | 4 | 5 | 6 | 6 | 7 | 7 | 7 | 7 | 8 |

#### Definition select_1_

Inverse to the **rank**_*σ*_ function, **select**_*σ*_ outputs the index of the *k*-th occurrence of symbol *σ* in the text.

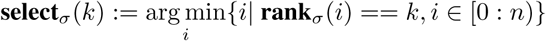

Example: **select**_1_ of *T* =`100101111010001` over Σ = {0, 1}

|  |  |  |  |  |  |  |  |  |  |  |  |  |  |  |  |
| --- | --- | --- | --- | --- | --- | --- | --- | --- | --- | --- | --- | --- | --- | --- | --- |
| $i$ | 0 | 1 | 2 | 3 | 4 | 5 | 6 | 7 | 8 | 9 | 10 | 11 | 12 | 13 | 14 |
| $T[i]$ | 1 | 0 | 0 | 1 | 0 | 1 | 1 | 1 | 1 | 0 | 1 | 0 | 0 | 0 | 1 |
| $k$ | 1 | | | 2 | | 3 | 4 | 5 | 6 | | 7 | | | | 8 |
| **select**<sub>1</sub>[$k$] | 0 | | | 3 | | 5 | 6 | 7 | 8 | | 10 | | | | 14 |

## 3. Algorithm

PriSeT operates in four steps:

1. **FM Index Step**. As a preprocessing, we compute the FM index on the complete corpus of all references. This needs only to be repeated when the references change.
2. **FM Frequency Step**. We compute the FM frequency for *k* in range [*κ*_min_ : *κ*_max_] and report all those *k*-mers exceeding a threshold F.
3. **Filter Step**. For each *k*-mer we check the first set of constraints *C_s_*, which must hold independently.
4. **Combine Step**. The remaining *k*-mers are combined reference-wise and checked for pair-wise fitting *C*_*p*_.

In order to handle the vast amount of *k*-mers and *k*-mer combinations, we use space efficient data structures and data types like bit encoding schemes for *k*-mers and references, and apply filters at early stages to reduce the number of *k*-mers before forming combinations.

### 3.1. FM Index Step

For indexing and *k*-mer report we use the recently published GenMap v1.0.1 by Pockrandt et al. (2019). As library input GenMap accepts a FASTA file or directory of FASTA files. Once computed, the FM index is used to answer frequency queries for arbitrary values of *k*. We modified GenMap to accept a frequency threshold parameter to avoid temporary storage of low frequent *k*-mers (see GenMap fork^8^). Algorithm 1 describes the call given GenMap’s interface.

#### Algorithm 1 FM index transformation of the library.

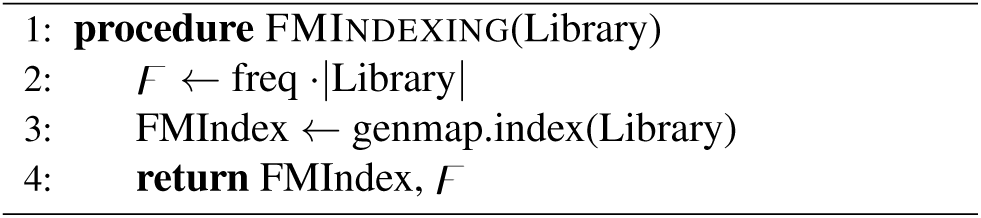

### 3.2. FM Frequency Step

Having computed the FM index over the library, we can now submit queries with varying values of *k*. GenMap’s frequency computation of *k*-mers is based on an algorithm described by Derrien et al. (2012), but introduces runtime improvements by cutting redundant searches (Pockrandt et al., 2019). GenMap provides location information of each *k*-mer by outputting the sequence identifier `SeqID` referring to the library sequence where a *k*-mer has been found and a relative position index `SeqPos`. These locations are collected in the `Locations` map (see Algorithm 2) with *k*-mers as keys and an occurrence vector as value.

#### Algorithm 2 *K*-mer frequency computation given FM index and frequency cutoff.

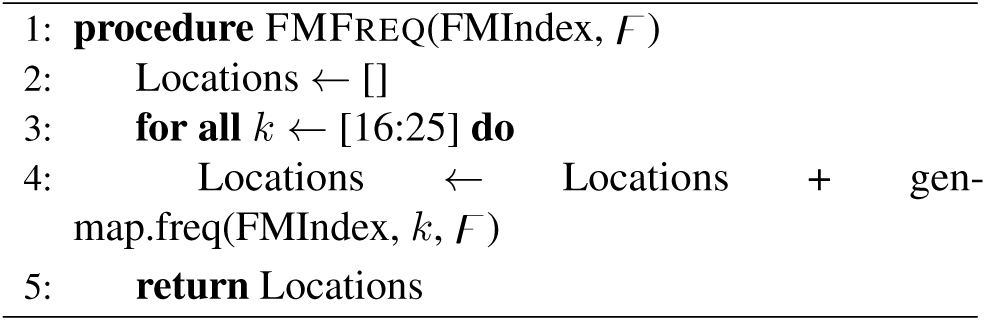

### 3.3. Filter Step

Each *k*-mer has to pass a chemical filter which checks molecular property constraints (see *C*_*s*_ in Table 2) that have to hold for single primers independent of possible matching partners. Applying the filter directly after the frequency step helps to reduce the number of *k*-mers by orders of magnitudes before running the combine procedure. The constraints *C*_*s*_ that are currently checked by PriSeT are amplicon length range, melting temperature, CG content, mono- or dinucleotide runs, and self-annealing patterns (see Table 2). amplicon ranges are constricted, because we first need to ensure that reads will overlap for the read merging step in a metabarcoding pipeline, and secondly, reads are usually trimmed to the same length – a precondition for most OTU clustering algorithms. The melting temperature in a PCR corresponds to the state where half of the DNA duplexes will be dissociated. Only when template DNA is single stranded, primer sequences can bind and the enzymatically driven elongation can take place. The temperature for the denaturation phase can be adjusted, but too high temperatures will lead to irreversible damage of the ingredients.

**Table 1.** List of symbols and their units or membership.

|  |  |  |
| --- | --- | --- |
| $F$ | absolute frequency cutoff for $k$-mer occurrences | $\mathbb{N}$ |
| $\kappa_{\min}$ | minimum $k$-mer length | nt |
| $\kappa_{\max}$ | maximum $k$-mer length | nt |
| $Tm_{\min}$ | minimum melting temperature | $^{\circ}\text{C}$ |
| $Tm_{\max}$ | maximum melting temperature | $^{\circ}\text{C}$ |
| $\tau_{\min}$ | minimum amplicon length | nt |
| $\tau_{\max}$ | maximum amplicon length | nt |
| $\Delta Tm$ | difference in melting temperatures | Kelvin |
| CG | Cytosine or Guanine occurrences | nt |
| AT | Adenine or Tyrosine occurrences | nt |

**Table 2.** Chemical Constraints for *k*-mers. The first set of constraints *C*_*s*_ applies to *k*-mers irrespective of their later orientation (forward or reverse). The second set of constraints *C*_*p*_ depends on the final orientation and the *k*-mer matching partner. *C*_*s*_ is applied in the **filter & transform** step and *C*_*p*_ checks in the **combine** step.

|  | Property | Recommendation |
| --- | --- | --- |
| $C_s$ | Primer Length | [16:25] |
|  | Read Length | [30:800] |
|  | Melting Temperature | [52:58] |
|  | CG content | [0.4:0.6] |
|  | Runs | not more than 4 same nt in a row |
|  | Dinucleotide Repeats | not more than 4 dinucleotides |
|  | Self-annealing | less than four consecutive nts and less than 50 % bond participation |
| $C_p$ | $\Delta T_m$ | 5 |
|  | AT tails | avoid (A/T)<sub>3</sub> tails |
|  | CG clamp | not more than 3 out of 5 nts at 3’ end are CG |
|  | Cross-annealing | less than four consecutive nts and less than 50 % bond participation |

Another criterion for primers being not too associative to the template is their CG content. Not only are `C`-`G` bonds twice as strong as `A`-`T` bonds, they are also more prone to mis-pairing. The recommended range is 40 - 60 % in proportion to their length. Mono- or dinucleotide runs are patterns where a single nucleotide or dinucleotide occurs consecutively repeated, i.e. *σ*_*x*_ or (*σ*_1_*σ*_2_)_*x*_ with *σ* ∈ Σ, *σ*_1_ ≠ *σ*_2_, and *x* ∈ 𝒩. These patterns tend to misprime. A maximal accepted number is four.

Self-annealing (or self-dimerization) occurs when copies of a primer form stable dimers. If their binding energy is relatively high, a significant amount will rather bind to primers than to the template and thereby deteriorating template amplification. The free Gibb’s energy is a measure for the bond strength. It numbers the energy that is released during bond formation and should not be below −6 Jmol^−1^. The amount depends on the positions and types of nucleotides involved, and can only be determined precisely via experiments. When looking at an alignment of two oligomers or primers, as a rule of a thumb, there should be not more than four annealing nucleotides in a row (connected annealing pattern) or not more than 50 % of the sequence (disconnected annealing pattern) involved in the bonding as shown in Figure 1.

**Figure 1.**
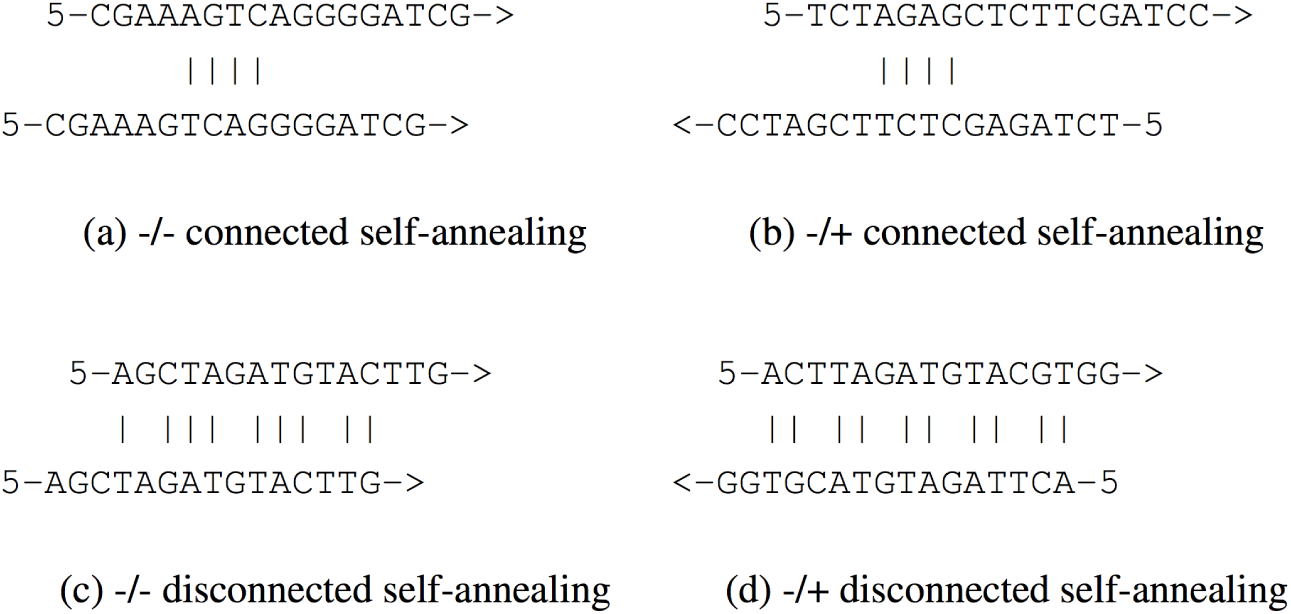
Critical annealing patterns. Oligomers in a) and b) have four self-annealing nucleotides in a row. In c) and d) more than half of the nucleotides participate in bonds. The oligomer orientation can be the same (-/-) or opposite (-/+).

The second set of constraints, *C*_*p*_, is used when evaluating primer pair candidates. One *k*-mer will be determined to be the forward primer on the minus strand from 5’ to 3’ direction, and the second *k*-mer the reverse primer on the plus strand. As mentioned earlier, denaturation and annealing phase temperatures can be adjusted, but we need to ensure that roughly the same amount of forward and reverse primers are dissociated to produce similar amounts of PCR products by allowing not more than five Kelvin differences in their melting temperatures. GC clamp refers to the amount of CG in the 5’-end of a primer. Especially at the 5’-end where the elongation takes place, a primer should bind stably, but not too strongly. Since `C` and `G` bind twice as strong as `A` or `T`, ideally one or two nucleotides should be `C` or `G`, but the last three should not be exclusively `A` or `T` (AT tail). For example, … `CGATA-3` and … `CGTCA-3` are not suitable, whereas … `AGTCA-3` is a suitable primer tail. Cross-annealing refers to the tendency that forward and reverse primer sequences form dimers (see Figure 1 b) and d)). The guidelines for avoidance are analogous to ones for self-annealing. The implementation of the constraint checks is described in more detail in the subsequent sections.

The initial amount of *k*-mers is quite large – up to 𝒪((*n* − *κ*_*min*_)(*κ*_*max*_ − *κ*_*min*_)) *k*-mers per reference sequence. For example, the Cryptophyta clade with a library size of 2 MB produces 3.8 million of *k*-mers^9^. We reduce space occupancy by not storing character strings, but a single unsigned 64 bit integer that holds multiple *k*-mers sharing the same prefix and location index. For quickly accessing *k*-mer positions, we use bit vectors in the length of the reference sequences. The next subsections describe in detail the encodings and how we apply the above described chemical constraints as filters on these encodings.

#### 3.3.1. Encoding of *K*-mers

When computing the *k*-mer frequency with varying values for *k*, one notices that for a specific position *i* in a reference *R*, it is likely that we will yield many *k*-mers of different lengths which successfully passed the frequency filter and start at the same index. In fact there are up to *κ*_*max*_ − *κ*_*min*_ + 1 many *k*-mers for the same location index. We introduce the `TKMerID` data type stored in a `uint64_t`, which primarily encodes the longest *k*-mer found at a specific text position (see Figure 3). We use the two bit encoding scheme shown in Table 3. The reduction to the four basic nucleotides is justified by the observation that substrings containing ambiguous nucleotides are meaningless, since a single mismatch between primer and template may lead to an amplification failure. The two-bit encoding scheme has the property that taking the complement of the integer produces its DNA complement. We use this property to implement a *k*-linear time annealing test.

**Table 3.**
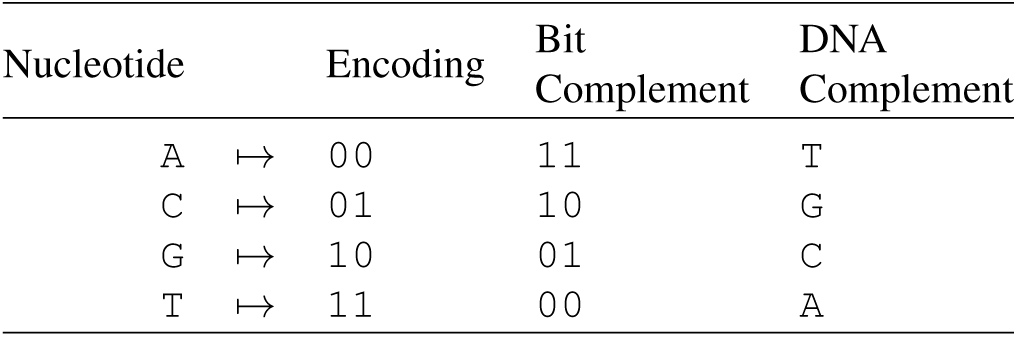
Two bit encoding of single nucleotides. Note that the bitwise complement operator produces the complement of an encoded nucleotide.

| Nucleotide |  | Encoding | Bit Complement | DNA Complement |
| --- | --- | --- | --- | --- |
| A | $\mapsto$ | 00 | 11 | T |
| C | $\mapsto$ | 01 | 10 | G |
| G | $\mapsto$ | 10 | 01 | C |
| T | $\mapsto$ | 11 | 00 | A |

**Figure 2.**
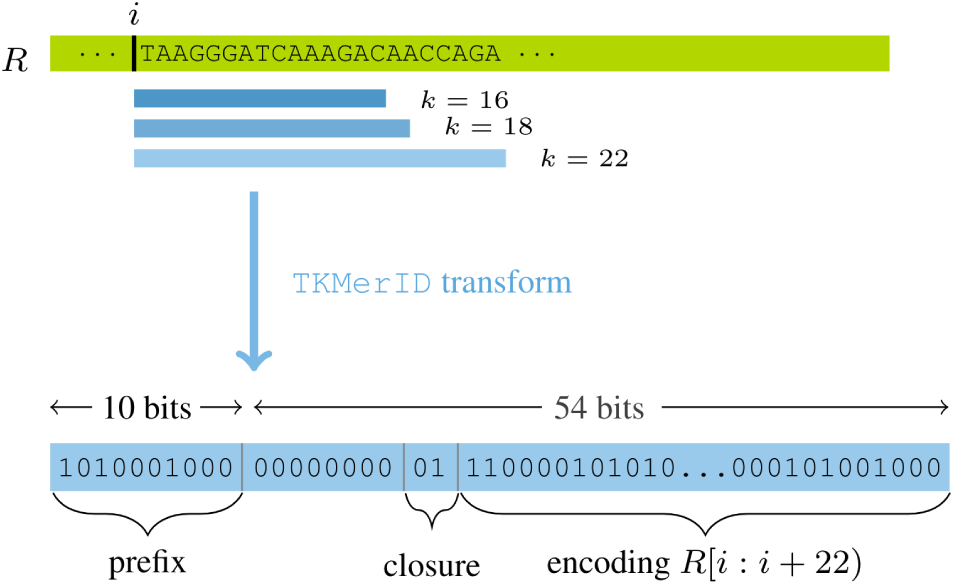
*K*-Mer encoding scheme. Top: All *k*-mers with the same start index also share the prefix. Redundancy is reduced by encoding the longest *k*-mer only. Bottom: In this example we encode the oligomer TAAGGGATCAAAGACAACCAGA with length bits set for 16, 18, and 22 in the prefix. The underlying data type is an `uint64_t`.

**Figure 3.**
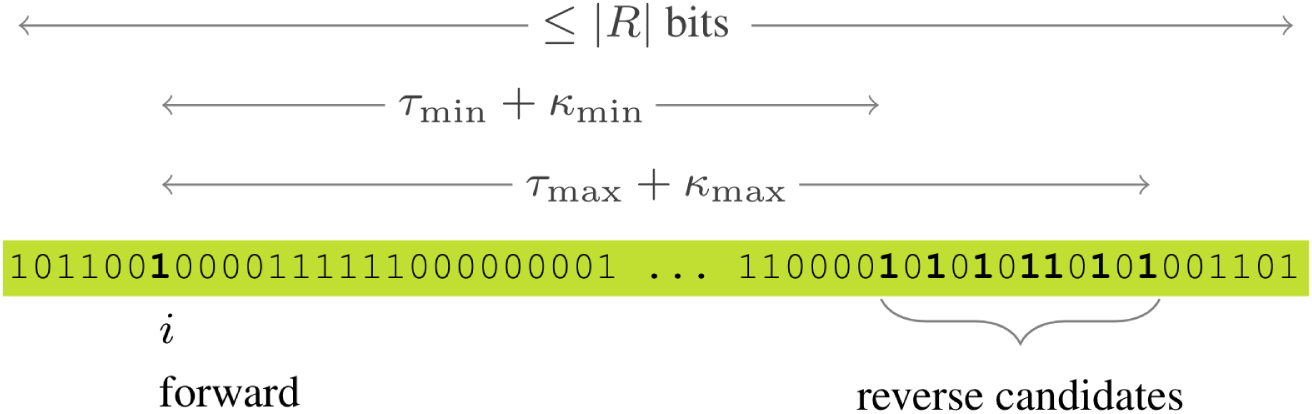
Encoding of TKMerID locations. Relative to a reference *R* PriSeT utilizes a `sdsl::bit_vector` with **rank**_1_ and **select**_1_ support in 𝒪(1). In this example the reference from which the bit vector originates from, has *k*-mers starting at the indices 0, 2, 3, 6, and so forth. Note, that in practice, we omit storing trailing zeros and the length of the bit vector corresponds to the position of the last *k*-mer occurrence. When searching for possible *k*-mer combinations to produce forward and reverse primer pairs, we have to parse for each forward `TKmerID` at a position *i* the window [*i* + *κ*_*min*_ + *τ*_*min*_ : *i* + *κ*_*max*_ + *τ*_*max*_].

The encoding consumes two bits per DNA base. Having at most *κ*_*max*_ = 25 nucleotides in a *k*-mer, 2 × 25 bits are consumed to store the longest accepted *k*-mer. We add two closure bits `01` to indicate the head of a *k*-mer. A prefix of `A`s (`00`_*x*_) would be ignored (or over-counted) otherwise. The at most 2 × 25 + 2 bits are stored in little-endian ordering in the suffix of a `TKMerID`. Two bits of the suffix will remain unused. The various lengths represented by a single `TKMerID` are stored in the preceding 10 bits. The *j*-th highest bit corresponds to *k* = *κ*_*min*_ + *j* − 1 (see Figure 3). Whenever we analyse the chemical properties of the encoded *k*-mers, we first mask out the prefix and then process the sequence starting with the lowest two bits. If a chemical constraint is not fulfilled for some *k*′, we only delete the prefix bit that corresponds to length *k*′. As long as the prefix is non-zero, we keep processing that `TKMerID` as a candidate primer.

#### 3.3.2. Encoding of *K*-Mer Locations

After the FM index computation, the original reference sequences are only needed once to lookup and encode frequent *k*-mers (see Filter Step). However, when combining *k*-mer candidates to form pairs, we need location information – because *k*-mers need to refer to the same reference and have to be in an offset range of [*τ*_*min*_ : *τ*_*max*_] nt.

We use two data structures per reference – a list to store the set of `TKMerID`s in order of occurrence and a *compact* data structure in form of a bit vector *B* in the length of the last *k*-mer occurrence. A set bit indicates the presence of a `TKMerID`. I.e., the *i*-th set bit in *B* corresponds to the *i*-th `TKMerID` in the KMerIDs list.

The *compact* data structure is augmented with **rank**_1_ and **select**_1_ support, which have 𝒪(1) runtime for queries^10^. Figure 3 illustrates the *k*-mer combination search over the transformed reference.

#### 3.3.3. Filter *C*_*s*_ for Single *K*-Mers

After having transformed the frequent *k*-mers into `TKMerID`s, each *k*-mer it represents needs to satisfy a set of constraints which must hold independently (see constraint set *C*_*s*_ in Table 2). All these constraint filters process `TKMerID`s linearly. For computing the melting temperature we use the Wallace rule which simply combines AT and CG counts linearly:

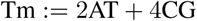

If for a length *k*_*l*_ the melting temperature exceeds Tm_max_, we can reset all length bits corresponding to lengths larger or equal to *k*_*l*_, because Tm grows monotonously with the length of the sequence. We count the nucleotides by comparing the last two bits with the single nucleotide codes (see Table 3), and then apply a right shift of two until we reach the closure bit.

The CG count can then directly be used to check for the CG content, which is recommended to be in the range of 40-60 %. However, since the CG content is relative to the sequence length, we have to check it for all encoded lengths of a `TKMerID`.

Patterns like mono- or dinucleotide runs are detected by comparing tailing bits of the code with the two-bit encoded patterns. For example, code to detect five consecutive cytosines:

~~~
uint64_t tail = (0b1111111111 & code);
bool has_run = (tail == 0b0101010101);
~~~

These patterns can be infixes of a *k*-mer. We therefore apply a right shift and comparison iteratively until 10 bits remain. In a similar fashion we proceed for dinucleotide runs.

With regard to our encodings, self-annealing can be determined by checking the shiftings of the original sequence against itself and its reversed copy (cross-annealing is analog). There are at most 2*κ*_max_ − 2·8 possible alignments and for each alignment we apply the ⊕ (XOR) operator to translate complementary nucleotides into `0b11` blocks. Critical self-annealing patterns then correspond to eight consecutive set bits starting at an even position, or an occurrence count of `0b11` exceeding *k/*2. In both cases, the presence can be detected in 𝒪(1) by counting bits in parallel and masking bits at indices of power of two in three steps.

Algorithm 3 puts together the previously described encodings of *k*-mers and their locations. `TKMerID`s need to be in order of occurrence, but *k*-mers are not processed in order. The strategy is to first set bits in the reference bit vector for *k*-mers, secondly to determine which values of *k* occur at a specific index, and thirdly, to fill the `TKMerID` vector.

### 3.4. *K*-mer Combiner

In the final combining step, we form pairs of two *k*-mers if they satisfy the second chemical constraint set *C*_*p*_ (see Table 2). We add **rank**_1_ and **select**_1_ support on every *B* ∈ ℬ to directly access single *k*-mer locations in 𝒪(1). When iterating over `TKMerID`s, **select**_1_ gives us its position index relative to the reference sequence. The search window of combinable reverse primers is then accessed by adding the amplicon length range. Again we apply **rank**_1_ on the window indices to address the associated `TKMerID`s in the list (see Algorithm 4). Pairs satisfying the matchability constraints *C*_*p*_ are output and can be ranked by coverage or barcode variation.

## 4. Experiments

We evaluated two scenarios for PriSeT. We applied PriSeT on an explicitly *uncurated* data set (data set for plankton) of short sequences (mostly 18S) taken from large clades to compute primer pairs exhibiting a large taxonomic coverage, and on complete viral genomes (SARS-CoV-2 genomes) to compute primer pairs producing species-distinctive amplicons. As a proof of concept test we additionally searched the primer result set on plankton data for primer pairs used in previous metabarcoding studies. At last, PriSeT’s runtime was analized on the plankton data set.

### Algorithm 3

Lookup and encoding of DNA *k*-mers and references. First the last *k*-mer occurrence for each reference is determined and a new bit vector instance resized accordingly. Each location L from the input map is associated with a concrete value for *k* and a list of occurrences represented as tuples of sequence identifiers and positions. The number of bit vector transformed references is referred to as *n*. In the first loop (line 4-6) bits are set in the bit vector for each *k*-mer occurrence. The second loop (line 8-12) composes the prefix by accumulating the length bits. The third loop (line 13-24) does the sequence lookup for the largest *k* occurring at a specific position.

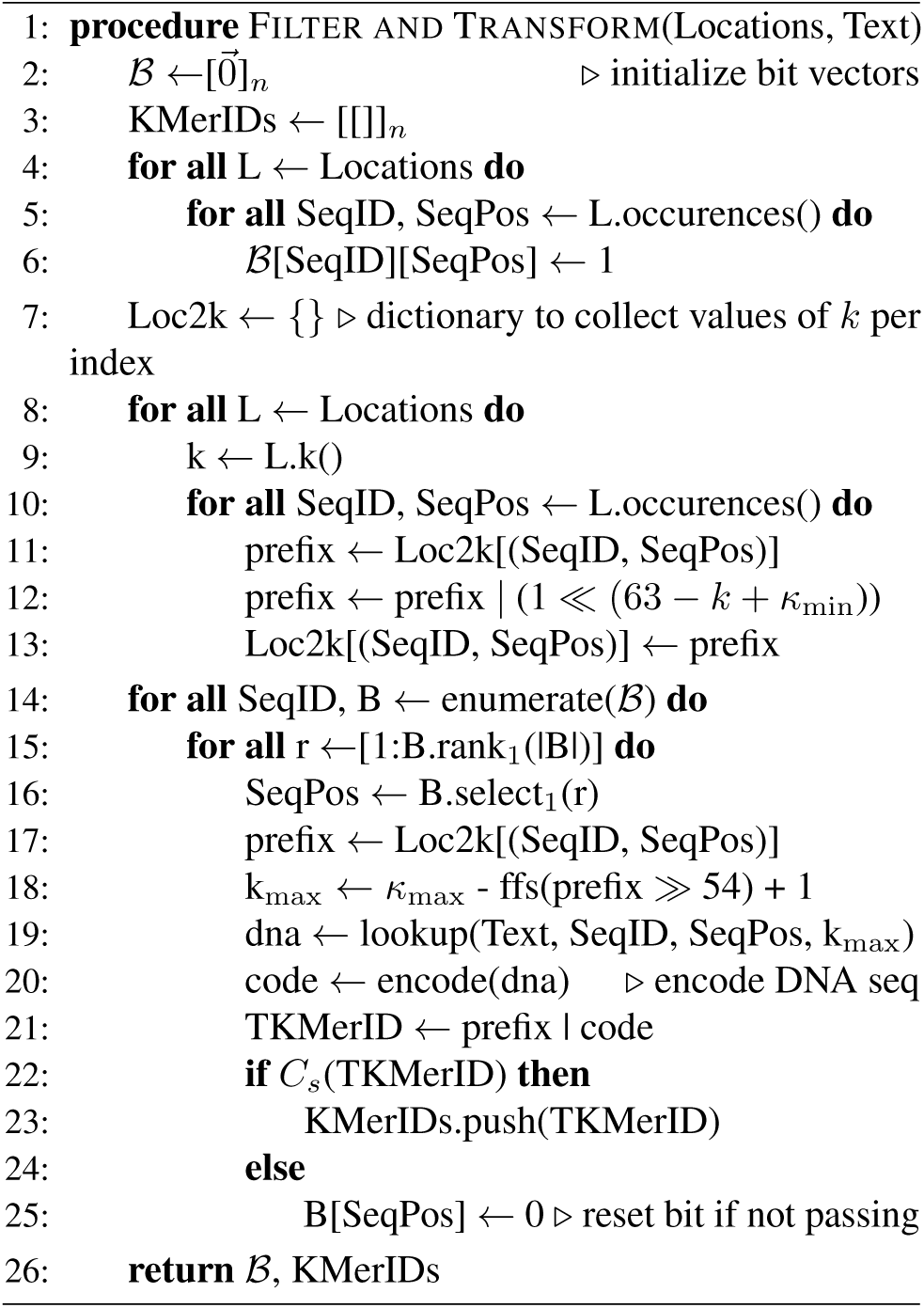

### Algorithm 4

Combine *k*-mers sequence-wise. For each position referring to a `TKMerID` in the bit vector transformed reference, we address candidates by adding an offset in the target amplicon length range. By using the **rank**_1_ on the window range, we only iterate over indices associated with `TKMerID`s. Then for each forward and reverse *k*-mer combination encoded by TKMerID1 and TKMerID2 (up to 100 combinations), we report chemically fitting pairs.

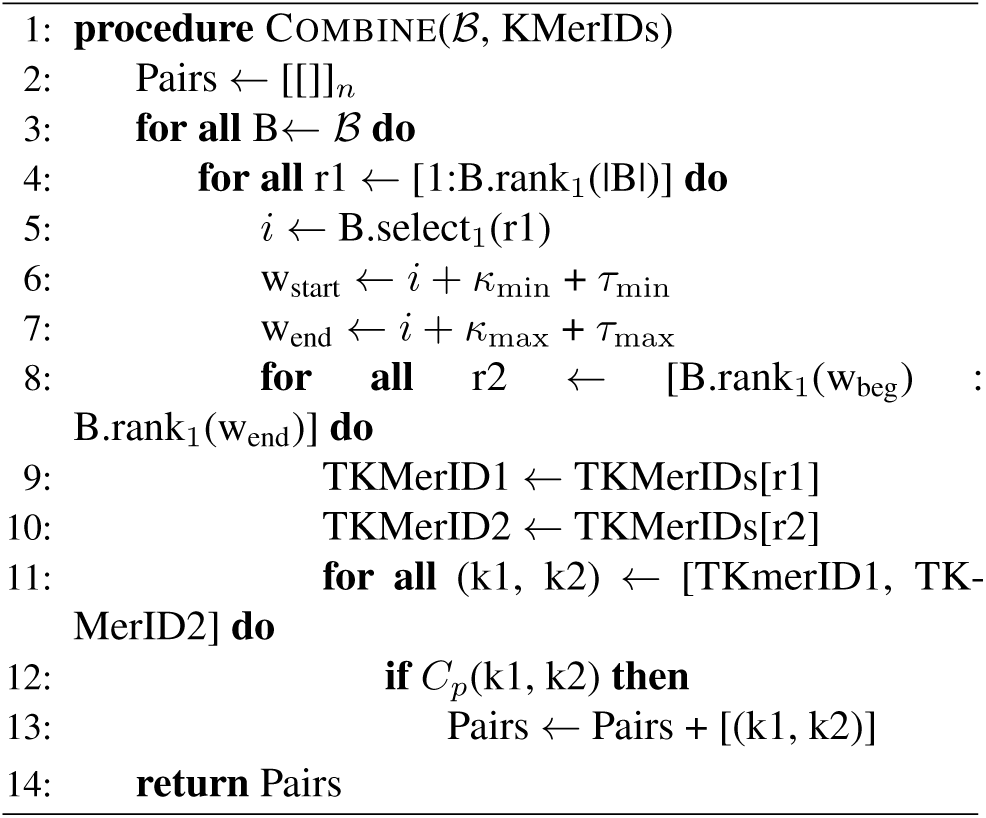

### 4.1. Data Set for Plankton

As a reference library, we sampled from NCBI GenBank’s *nt* dataset (Benson et al., 2012)^11^, which contains non-human sequences from various sources. The prevalent sequence length range is between 400 to 2500 bases. We picked 19 clades that include Eukarya typically found in freshwater plankton samples ranging from phyto-to zooplankton and fungi. For each taxon within a clade that contained at least one accession assigned to it, we sampled at most three accessions to remove the sequence bias introduced by highly populated taxa. Table 4 lists the clades, the number of taxa, taxa with at least one accession (Covered), the total number of accessions and the library size in megabytes (MB). Between 1.38 (Rotifera) to 2.5 (Charophyceae) accessions were sampled from covered nodes, which indicates the sparse population of the reference library.

**Table 4.**
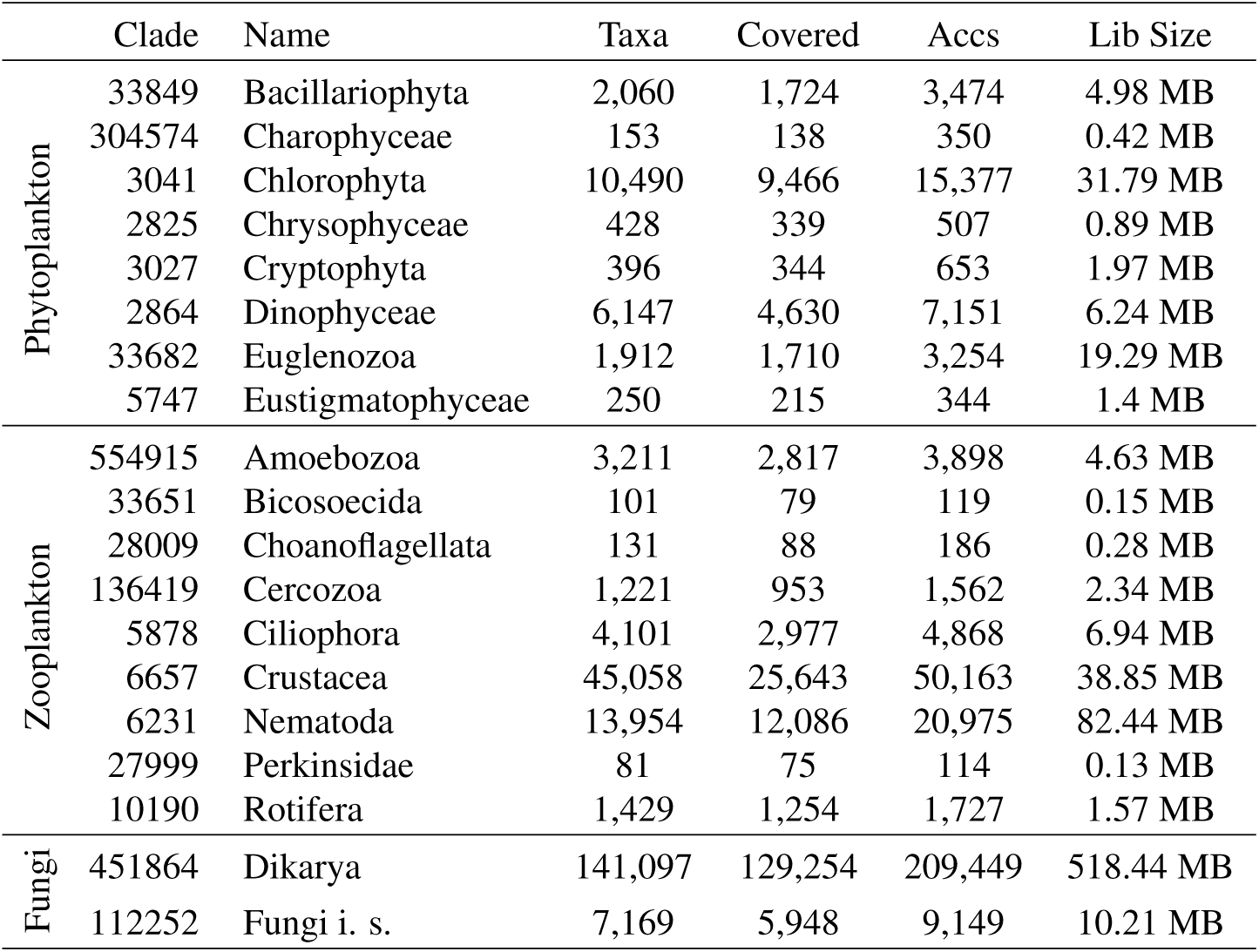
Data set used for the primer verification test, the *de novo* search, and the runtime analysis. Taxonomic identifiers in the clade column follow the NCBI nomenclatura, Taxa refers to the total number of nodes (including virtual ancestors), Covered the number of taxa having at least one accession assigned to it, Accs to the total number of collected accessions and Lib Size to the size of the fasta file containing all accessions.

### 4.2. Verifying Established Primer Pairs for Plankton

Bacillariophyta (diatoms) and green algae (mostly Chlorophyta) are among the most abundant organisms in freshwater and marine plankton communities. Bacillariophyta are small (2 - 200 *µ*m) and their characteristic silica cell walls allow morphological identification by trained experts. Their role in ecosystem function and for environmental monitoring and water quality assessment cannot be underestimated – they contribute approximately 20 % of global oxygen production and represent nearly half of the organic material in the oceans. The DIV4 primer pair (see Table 5) was specifically designed for Bacillariophyta by Visco et al. (2015) with an expected amplicon length of ∼280 nt. Hadziavdic et al. (2014) optimized primers towards a large Eukaryota coverage (universal eukaryotic primers) by aligning all sequences from the SILVA database and computing the entropy at each alignment position. From the regions with low entropy they identified eight forward, and six reverse primer candidates. Here we use F-566a as forward, and R-1200 as reverse primer (E14 hereafter).

**Table 5.**
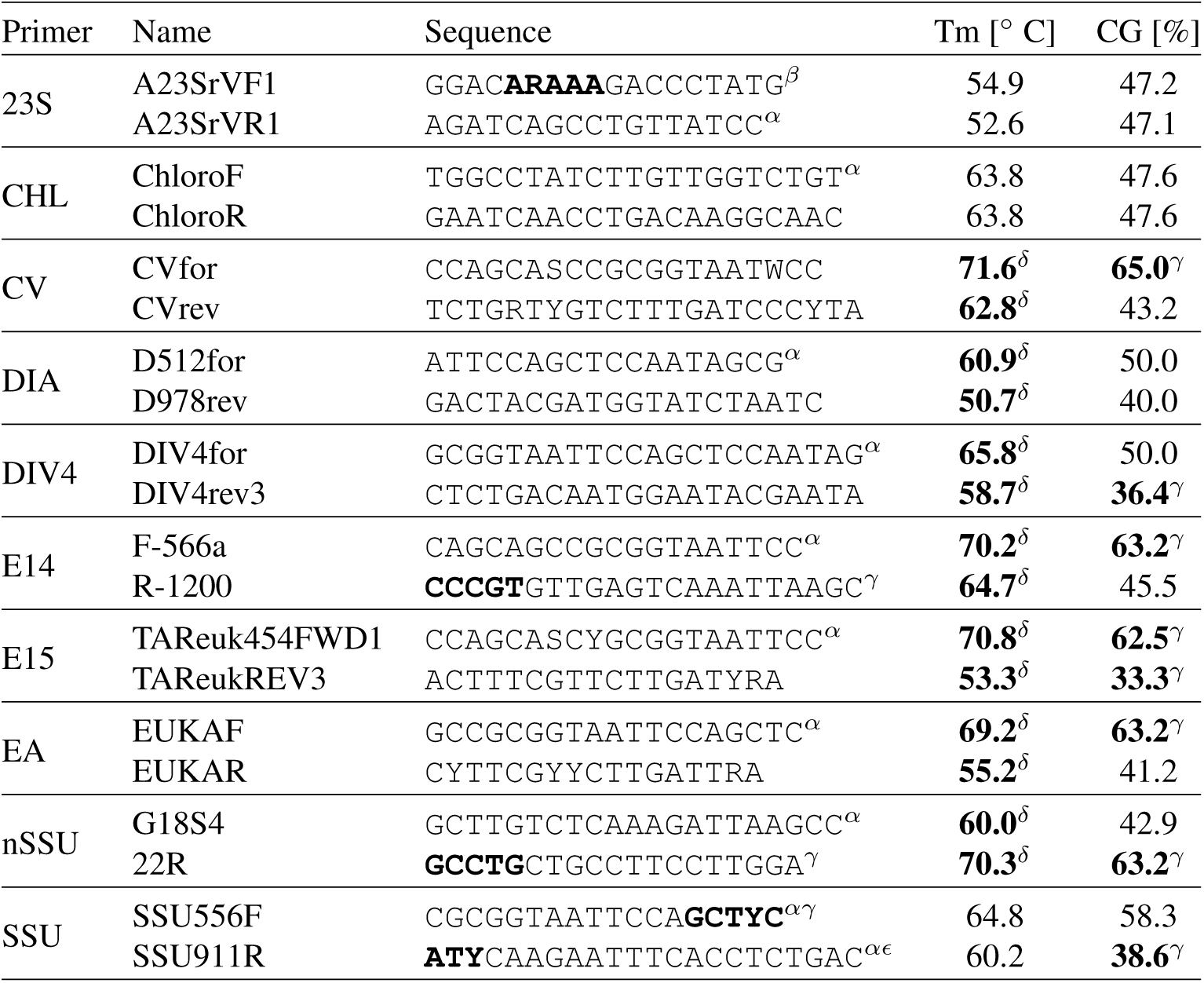
Selected primer sequences for the verification experiment. Sequences are noted in 5’ to 3’ direction. Melting temperatures are computed using a modified nearest-neighbor method described by Breslauer et al. (1986) assuming a primer concentration of 0.5 *µ*m and salt 50 mM. For ambiguous encodings, the average of both extrema is computed. Sequence parameters violating primer design recommendations are written bold. Critical structures are marked: self-annealing (*α*), mononucleotid runs (*β*), CG clamps (*γ*), exceeding ΔTm (*δ*), (A|T)_3_ tails (*ϵ*), and exceeding CG content ranges (*γ*).

Other tested primers are 23S by Yoon et al. (2016) developed for marine phytoplankton (23S hereafter), ChloroF/R by Moro et al. (2009) for Chlorophyceae and Bacillariophyceae (CHL hereafter), CVfor/rev by Boscaro et al. (2017) for freshwater ciliates (CV hereafter), D512for/D978rev by Zimmermann et al. (2011) for diatoms (DIA hereafter), TAReuk454FWD1/ TAReukREV3 by Stoeck et al. (2010) for the V9 region of marine Eukaryota, EUKAF/R by Moreno et al. (2018) for the 18S region of Protozoa (EA hereafter), G18S4/22R by Blaxter et al. (1998) for Nematoda (nSSU hereafter), and SSU556F/SSU911R by Kirsty et al. (2017) for Dinoflagellata (SSU hereafter). Some of these primers were designed with respect to a specific organism group, however, it is expected that they are also effective in other clades.

Table 5 lists the 10 selected primer pairs targeting 18S that we searched for in the reference library. It is remarkable that not a single pair has chemically optimal properties. At least one sequence of a pair shows a self-annealing pattern, seven pairs differ significantly in melting temperatures (independent of computation method, i.e. Wallace rule or nearest-neighbour method), three sequences have CG clamps at their 3’ ends, eight sequences have exceeding CG contents, 23S forward contains a run of five adenine bases (`R` substitutes `A` or `G`), and SSU911R has an (A|T)_3_ tail. We therefore relaxed the chemical constraints for the verification experiment by allowing a larger melting temperature range and difference in ΔTm, a larger CG content range, and we deactivated the self-annealing filter (see column ‘Verification’ in Table 6).

**Table 6.**
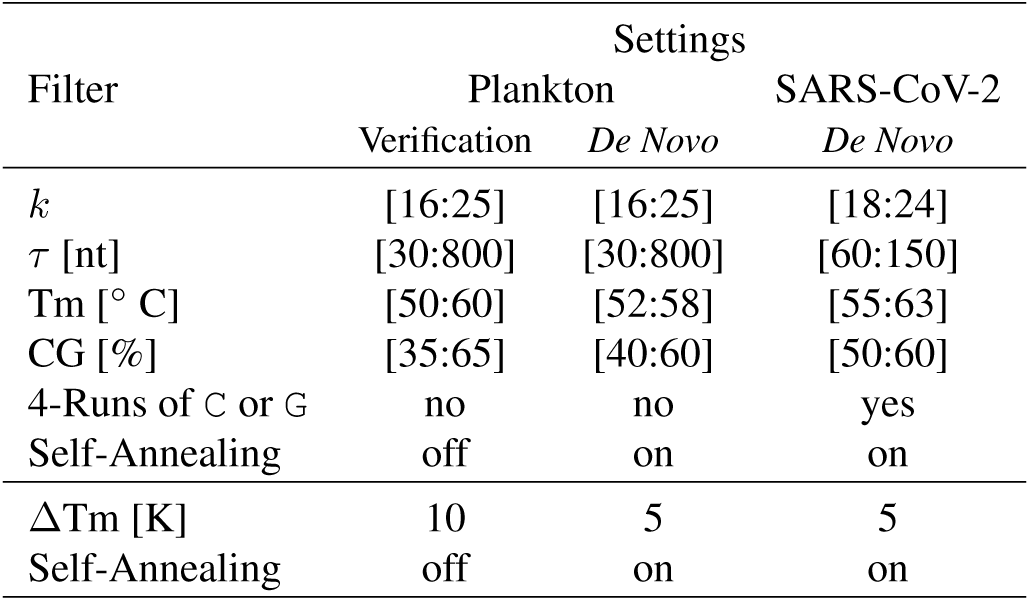
Filter Settings in PriSeT. Unlisted properties follow the recommended ones in Table 2.

Given the sampled library, we first computed the ground truth by searching directly for the known primer sequences. We used a simple linear text search and marked all data sets having at least one accession with forward and reverse primers matching as indicated by a presence (1) or absence (0) bit in the denominator of each cell in Table 7. We then ran PriSeT with the relaxed constraints listed in the Verification column of Table 6 and searched its result set for the published primers listed in Table 5. We indicated their presence or absence in the nominator.

**Table 7.** Primer presence test for established primer pairs. *x/y* refers to found by PriSeT (*x* = 1) or not (*x* = 0) versus found in the library (*y* = 1) or not (*y* = 0). Primers not discovered by PriSeT did not pass either the first chemical filter set *C*_*s*_ (labeled with ∗) or the pair filter set *C*_*p*_ (labeled with ∗∗).

|  | Clade | 23S | CHL | CV* | DIA | DIV4* | E14 | E15** | EA** | nSSU | SSU |
| --- | --- | --- | --- | --- | --- | --- | --- | --- | --- | --- | --- |
| Phytoplankton | 33849 | 1/1 | 0/0 | 0/0 | 1/1 | 0/1 | 1/1 | 0/1 | 0/1 | 1/1 | 1/1 |
|  | 304574 | 1/1 | 0/0 | 0/0 | 0/0 | 0/0 | 1/1 | 0/1 | 0/1 | 0/0 | 1/1 |
|  | 3041 | 1/1 | 1/1 | 0/0 | 1/1 | 0/0 | 1/1 | 0/1 | 0/1 | 1/1 | 1/1 |
|  | 2825 | 0/0 | 0/0 | 0/0 | 1/1 | 0/1 | 1/1 | 0/1 | 0/1 | 1/1 | 1/1 |
|  | 3027 | 1/1 | 0/0 | 0/0 | 0/0 | 0/0 | 1/1 | 0/1 | 0/1 | 1/1 | 1/1 |
|  | 2864 | 1/1 | 0/0 | 0/1 | 1/1 | 0/1 | 1/1 | 0/1 | 0/1 | 1/1 | 1/1 |
|  | 33682 | 1/1 | 0/0 | 0/0 | 0/0 | 0/0 | 1/1 | 0/1 | 0/0 | 0/0 | 0/0 |
|  | 5747 | 1/1 | 1/1 | 0/0 | 1/1 | 0/0 | 1/1 | 0/1 | 0/1 | 1/1 | 1/1 |
| Zooplankton | 554915 | 0/0 | 0/0 | 0/1 | 0/0 | 0/0 | 1/1 | 0/1 | 0/1 | 1/1 | 1/1 |
|  | 33651 | 0/0 | 0/0 | 0/0 | 0/0 | 0/0 | 1/1 | 0/1 | 0/1 | 1/1 | 1/1 |
|  | 28009 | 0/0 | 0/0 | 0/0 | 0/0 | 0/0 | 1/1 | 0/1 | 0/1 | 1/1 | 1/1 |
|  | 136419 | 0/0 | 0/0 | 0/0 | 0/0 | 0/0 | 1/1 | 0/1 | 0/1 | 1/1 | 1/1 |
|  | 5878 | 0/0 | 1/1 | 0/1 | 0/0 | 0/0 | 1/1 | 0/1 | 0/1 | 1/1 | 1/1 |
|  | 6657 | 0/0 | 0/0 | 0/0 | 0/0 | 0/0 | 1/1 | 0/1 | 0/1 | 1/1 | 1/1 |
|  | 6231 | 0/0 | 0/0 | 0/0 | 0/0 | 0/0 | 1/1 | 0/1 | 0/1 | 1/1 | 0/0 |
|  | 27999 | 0/0 | 0/0 | 0/0 | 0/0 | 0/0 | 1/1 | 0/1 | 0/1 | 0/0 | 1/1 |
|  | 10190 | 0/0 | 0/0 | 0/0 | 0/0 | 0/0 | 1/1 | 0/1 | 0/1 | 0/0 | 0/0 |
| Fungi | 451864 | 0/0 | 0/0 | 0/1 | 0/0 | 0/1 | 1/1 | 0/1 | 0/1 | 1/1 | 1/1 |
|  | 112252 | 0/0 | 0/0 | 0/1 | 1/1 | 0/0 | 1/1 | 0/1 | 0/1 | 1/1 | 1/1 |

We could recover all those primer sequences that satisfied the relaxed constraints. The forward primer of DIV4, and both primers of CV did not pass the relaxed Tm filter (above 65° C) and failed therefore for *C*_*s*_, and the primer pairs E15 and EA failed for *C*_*p*_ because of their excessive differences for Tm (more than 8 Kelvin).

### 4.3. *De Novo* Computation for Plankton

To yield chemically uncritical primer pairs, we reset the constraints to the recommended ranges (see column ‘De Novo’ of Table 6) and reported the 50 most frequent *k*-mer pairs for each clade (labeled with hash values computed from forward and reverse sequence). We additionally computed frequency, coverage, and variation for the published primers (Table 5) via text search without that they underwent PriSeT’s filtering.

The *de novo* and published primer result sets were either ranked by taxon coverage (columns 2-4 in Table 8) or by amplicon variation (columns 5-7 in Table 8). From the *de novo* set and the published primers set, we reported only the highest ranked ones. For example, for Charophyceae (clade 304574), the highest ranked primer by coverage is `d8d47dc9b873d02b` – a *de novo* computed primer, and the highest ranked published primer is EUK14. In the joint set, however, there are more *de novo* primer pairs that have a higher rank than EUK14, but are not reported here for the sake of briefty. When ranking by coverage, for 11 out of 19 clades a *de novo* primer outperformed all published primers. Since coverage optimizes towards broadness and variation towards amplicon distinguishability, we expect that the top primers for the two ranking methods may differ. In 14 out of 38 cases the top primers of the *de novo* or published sets are identical. In 24 cases higher variation is paid with lower coverage and *vice versa*. When ranking by number of unique amplicons (variation), we could identify seven *de novo* primers that performed equally well (see Table 8).

**Table 8.**
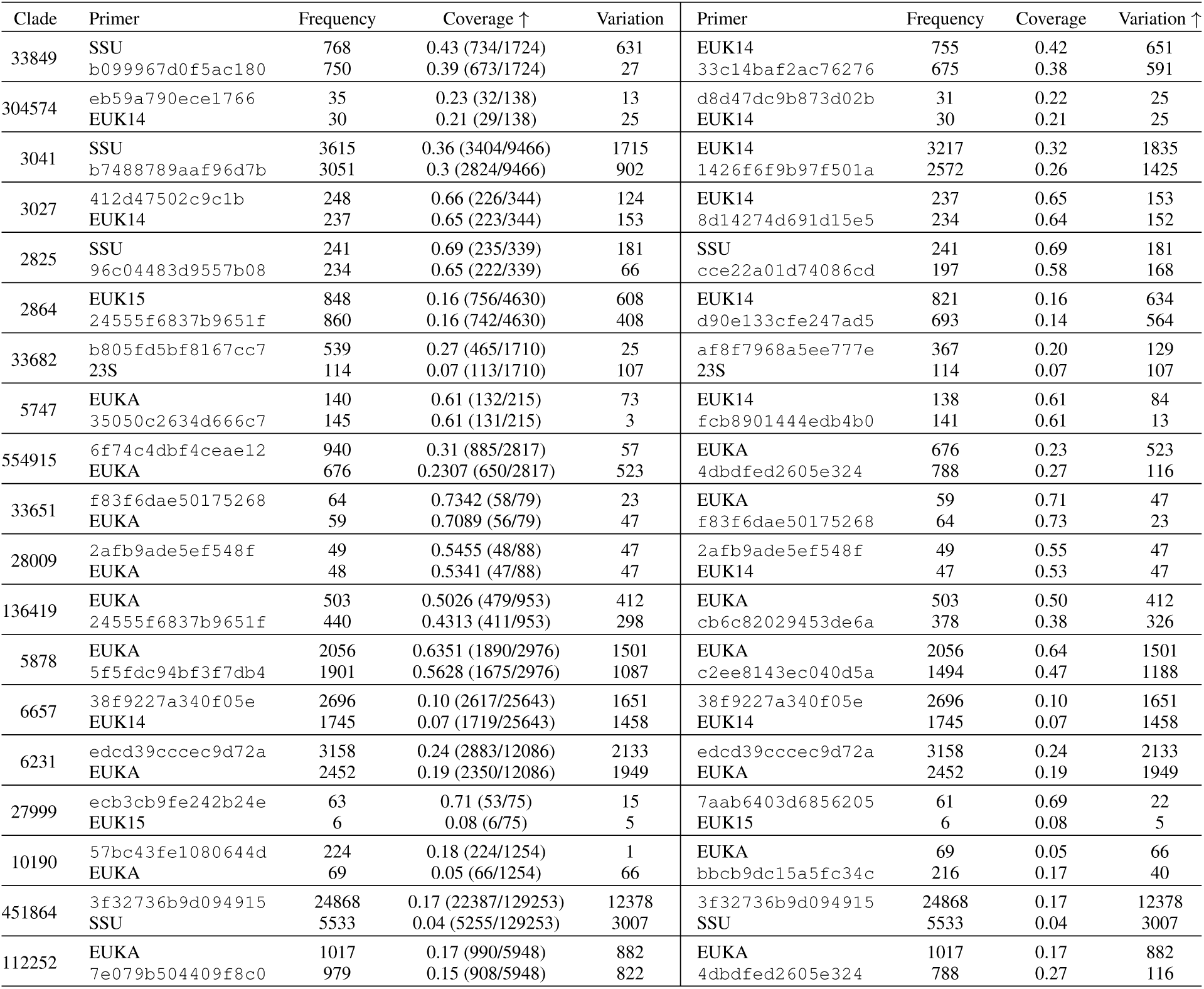
*De novo* computation and statistical comparison with published primers ranked by Coverage (left) or Variation (right). Of both sets the top performing primer pair in terms of coverage or variation were picked. Frequency reflects the raw number of occurrences in the clade, whereas Coverage computes how many taxa of each clade were covered in proportion to taxa with references, and Variation the total number of amplicons. Note, that PriSeT computed primer pairs with narrowed constraints – primer pairs like SSU, EUKA, etc., would not emerge in the result set (see Table 6). When ranking by coverage, for 11 out of 19 clades PriSeT identifies at least one new primer pair with a higher coverage rate than the published primers. Whereas when ranking by amplicon variation for seven out of 19 clades PriSeT found at least one more or equally performant primer pair.

In spite of the reference sparsity and sequential diversity – references not only come from 18S, but other genomic regions, PriSeT could identify for some clades new primer pairs that fit chemically while having a greater coverage or variation. Because of the known problem of data sparseness for plankton clades and the obvious fact that GenBank’s reference sequences stem from PCR amplicons with already published primer sets, we had lower expectations.

Published primers outperform *de novo* primers by at most 7 %, and *de novo* the published ones by at most 70 % for coverage. However, there may exist other performant primer pairs for specific clades that we are not aware of. We chose the most promising ones found in the PR2 database^12^. We also tested EUKA/B from Medlin et al. (1988), Cerc479F/Cerc750R by Harder et al. (2016), and DimA/DimB by Cannon et al. (2018), which either did not show up or they occurred with very low frequencies.

From the published primer set EUK14 and EUKA seem to be very versatile primer pairs. However, they do not satisfy *C*_*s*_ or *C*_*p*_ and might require more experience when applied in a PCR. The complete list of top performing primers is available on GitHub^13^.

### 4.4. Dataset for SARS-CoV-2

We selected 19 complete SARS-CoV-2 genomes^14^ from the subgenus Sarbecovirus for PriSeT to compute primer pairs satisfying the RT-PCR constrains listed in Table 6. The single genome size was roughly 30,000 bp. In order to filter for primer pairs producing amplicons with no occurrences in other Orthocoronavirinae genomes, we selected all available genomes from GenBank – 24 Alphacoronavirus, 5 Betacoronavirus (excluding *Sarbecovirus*), and 2 Gammacoronavirus genomes (see Figure 4).

**Figure 4.**
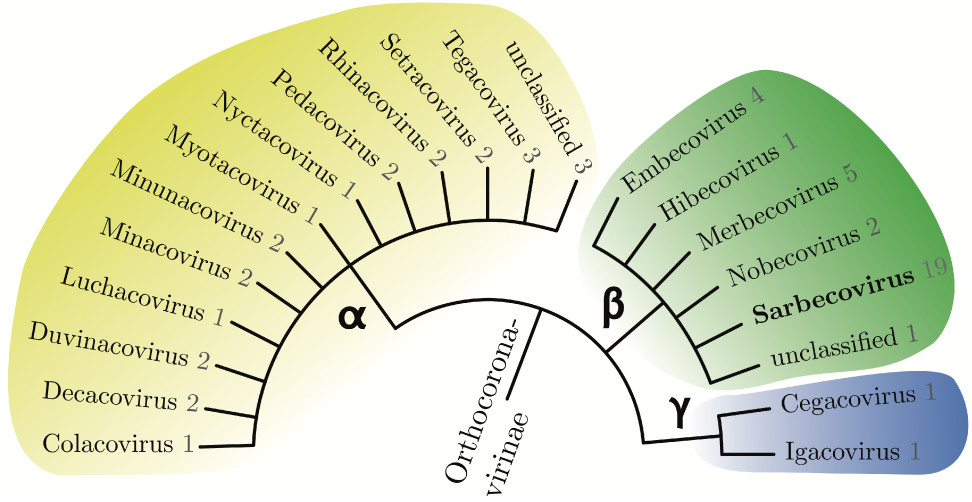
Taxonomy of Orthocoronavirinae to subgenus level. The numbers indicate how many complete genomes were used for the primer *de novo* search of each subgenus. From the *Sarbecovirus* subgenus all 19 genomes are assigned to the species SARS-CoV-2. Their proximity to Bat SARSr-CoV (another *Sarbecovirus*) was observed in the amplicon analysis (see experimental section.

### 4.5. *De Novo* Computation for SARS-CoV-2

PriSeT produced 286 primer pairs. We subsequently filtered for those producing sequences that cannot be found in one of the other 39 coronavirus genomes based on a 100 % sequence identity. For the 114 remaining primer pairs, we ensured amplicon distinction by launching two BLAST queries for each of the 114 amplicons against GenBank’s nucleotide collection (nt/nr) online^15^. The first query was run on the complete nt/nr data set and the second on the complete nt/nr data set except SARS-CoV-2 (taxid 2697049) to ensure that we did not miss relevant matches with non SARS-CoV-2 entries.

None of the primer pairs produced amplicons with 100 % identity and 100 % coverage for non *Sarbecoviruses*. Of the 114 primer pairs 5 pairs had 100 % sequence identity with a single accession of a *Sarbecovirus* isolated from the pangolin, but no other relevant matches. There were 109 primer pairs producing amplicons with no other occurrences. Out of the 109 primer pairs, there were 12 pairs with amplicons that were distant from even closely related viruses, i.e. sequence identity < 97 %, and 97 had a proximity (but not identity) to at most two other accessions, namely, two *Sarbecovirus* species isolated from a bat and the pangolin. The first one is associated with the recent pneumonia outbreak (Zhou et al., 2020).

The complete list of primer sequences, amplicons, and comments about closest matches can be found in the supplements (`primer_priset_covid19.csv`). The approximate amplicon locations relative to the genome are shown in Figure 5. For the sake of brevity only the first four digits of the primer identifier (first column in corresponding file) is noted down.

**Figure 5.**
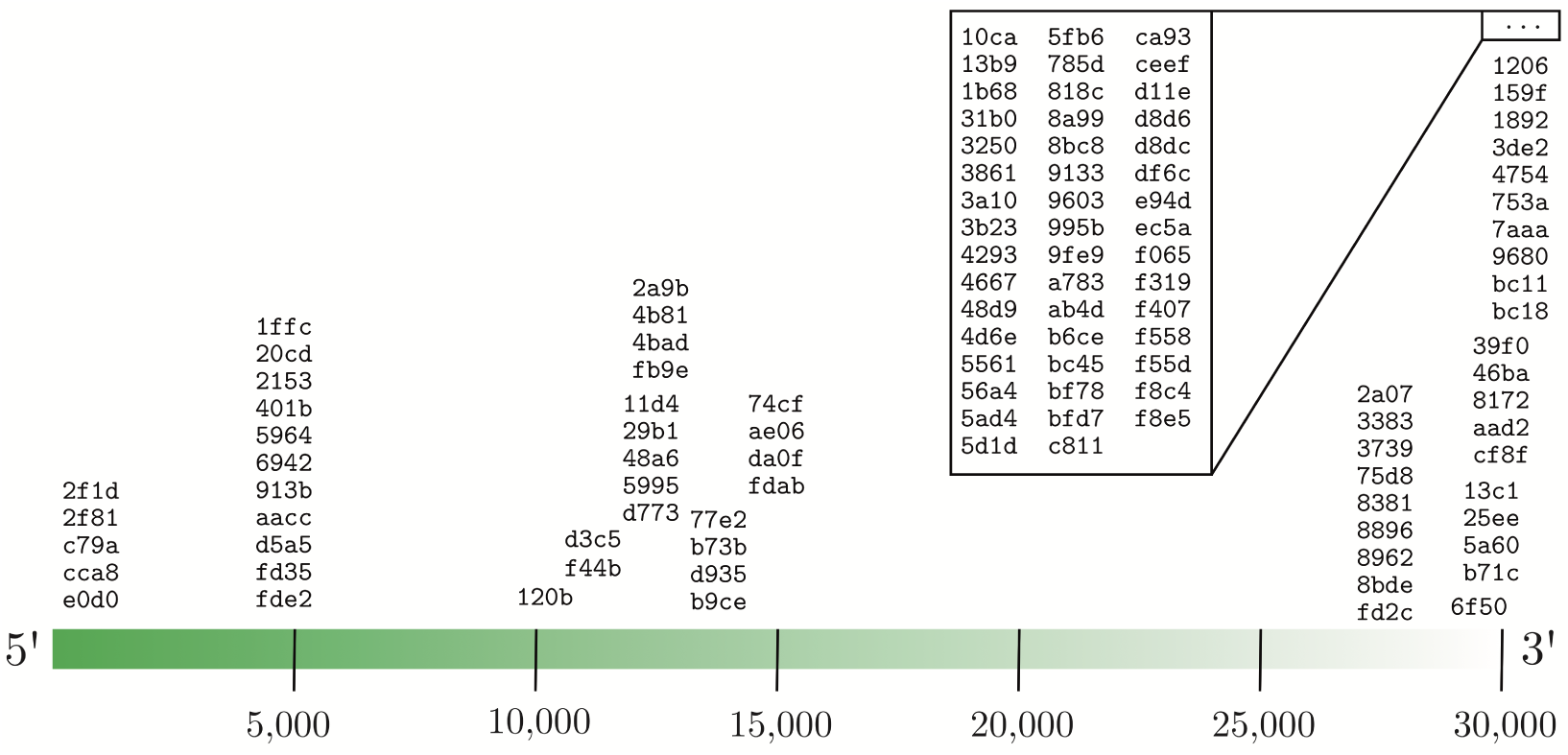
Approximate amplicon locations of *de novo* computed primer pairs for SARS-CoV-2. The amplicon identifiers are the first four digits of the primer pair identifiers in the supplemental file primer_priset_SARS-CoV-2.csv.

### 4.6. Performance on Plankton Data Set

For each clade we measured the step-wise runtimes and the total runtime excluding the FM index computation (see Figure 6). The index computation needs to be done only once and upon updates of the original library. For our data set, the FM index consumed about 4.2 times more space than the library it is computed on.

**Figure 6.**
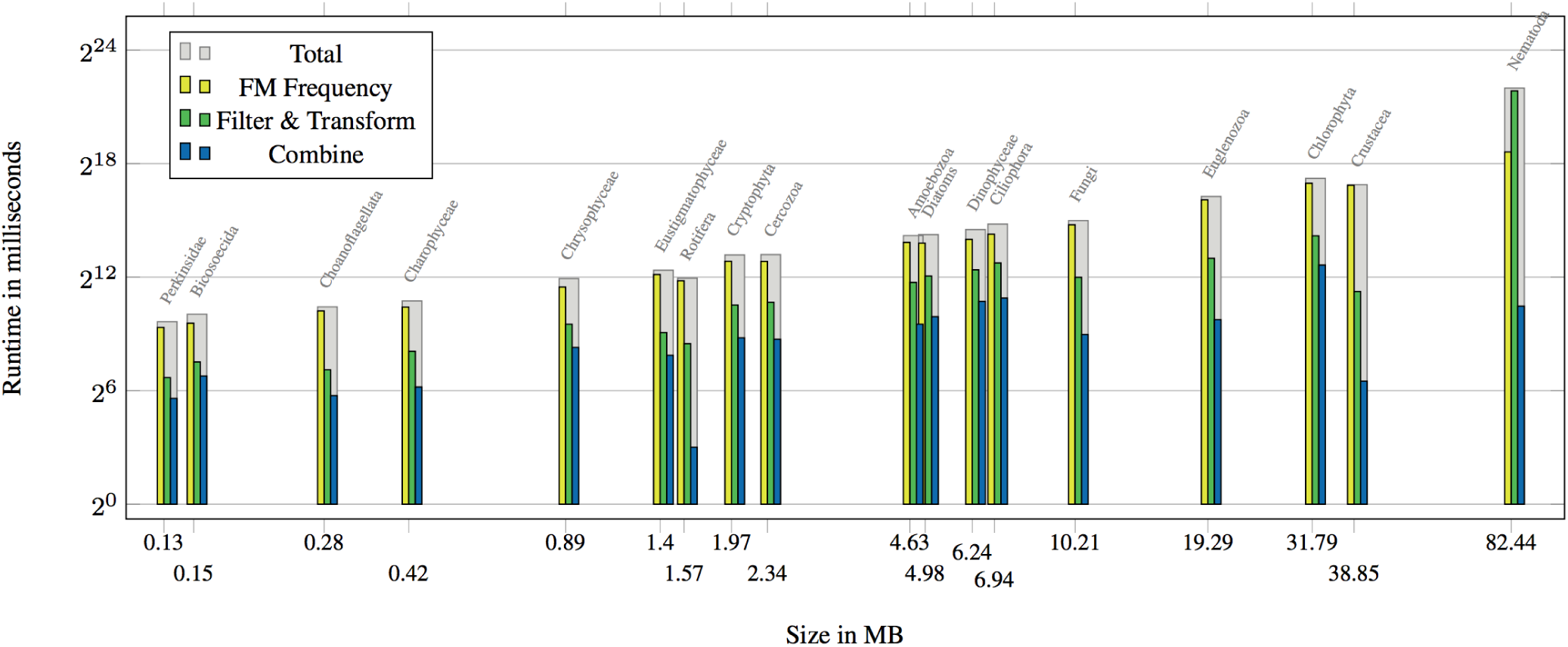
Runtimes for all clade data sets broken down to frequency computation, filter & transform, and combine step. The *k*-mer frequency cutoff was set to 5% w.r.t. the number of references per clade. The results for Fungi (clade 451864, 500 MB large) are omitted here due to the necessity of setting the cutoff to 10%, which results in runtimes comparable with clade 6231 (82 MB). Both axes are log_2_-scaled.

We set the relative frequency cutoff to 5%. The number of *k*-mer locations grows therefore linearly with the library size. We have chosen not to show the performance on a synthetic data set, because *k*-mer frequencies and dropout rates are defined by inherent sequence properties (entropy, repeats, etc.), which are obviously not homogeneous over all clades. A single clade or synthetic data set would produce non-representative, and even misleading results. For example clade 6657 has a library being 7 MB larger than the one of clade 3041. Surprisingly, clade 6657 produces only 4.83 million *k*-mers, compared to 43,9 million *k*-mers of clade 3041 (see Figure 7). For the largest data set (clade 451864 of Dikarya), we had to raise the frequency cutoff to 10 %, for not running into memory issues caused by the vast amount of *k*-mers (see Discussion 5).

**Figure 7.**
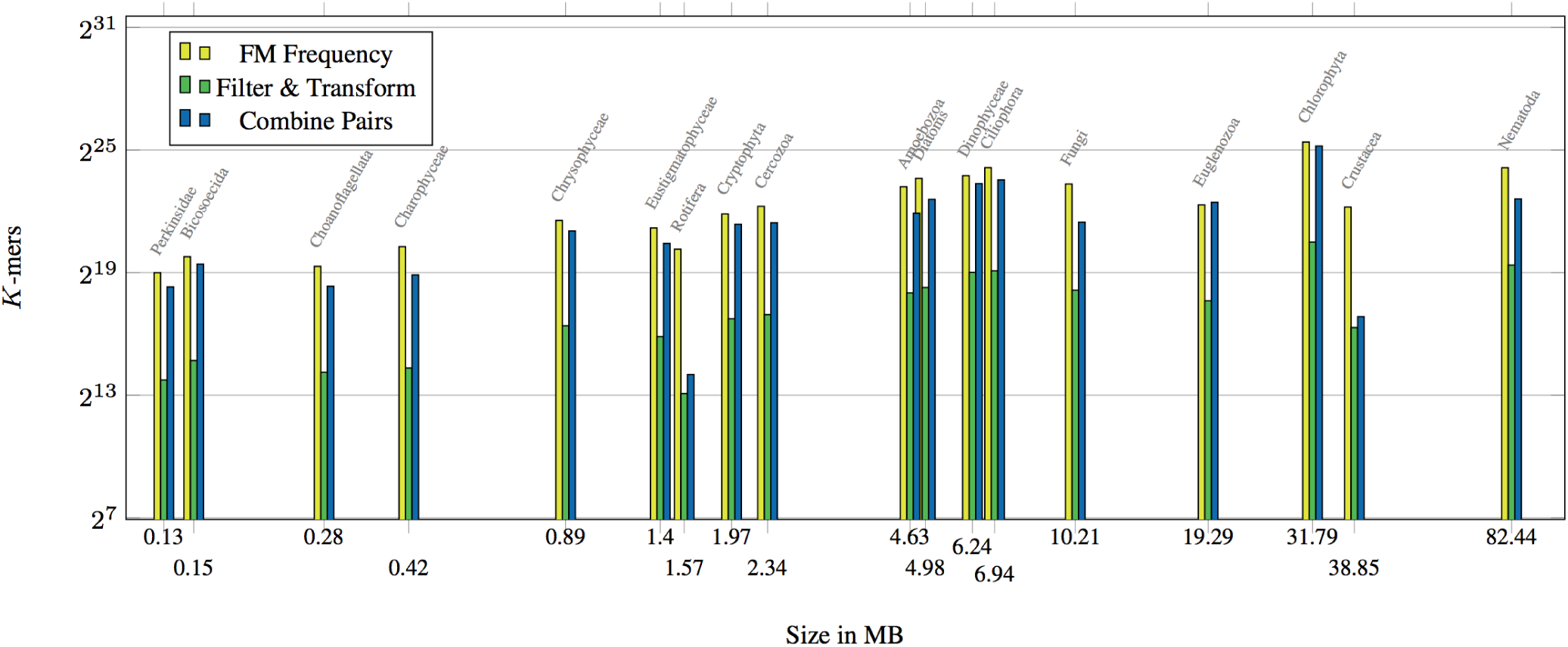
*K*-mer Counts for all clade data sets broken down to frequency computation, filter & transform, and combine step. *K*-Mers for all clade data sets counted after the **frequency** computation, **filter & transform**, and **combine** steps. For the **combine** step pairs are counted, not *k*-mers. The settings are the same as for Figure 6.

We sorted the data sets by size (abscissa) and log_2_-scaled abscissa and ordinate for readability. The total runtime for the smallest data set Perkinsidae (clade 27999 with 0.13 MB) is ≤ 1 second, for Fungi (clade 112252 with 10.21 MB) 33 seconds, and for a large dataset like Nematoda (clade 6231 with 82.55 MB) 70 min.

The **frequency** computation contributes the most to the total runtime. This is due to the large number of possible *k*-mers within a library. After the **filter & transform** step the number of *k*-mers is reduced drastically, s.t. the expensive **combine** step with a runtime in the size of all *k*-mer positions times the window size remains relatively low.

The *k*-mer dropout rate during **filter & transform** and **combine** is highly dependent on the sequence structure within the clades as indicated by the non-linearity of **filter & transform** and **combine** runtimes in proportion to the original library size.

Theoretical upper limits can be seen in Table 9. The **frequency** computation performs a single *k*-mer look-up in *O*(*k*) where *k* is the length of the *k*-mer to be looked up. Additionally, all occurrence locations need to be gathered, which depend on the number of occurrences *occ* and the total library length *N*. This can be done in *O*(*k* + *occ*) by exploiting the lexicographical ordering of the index (Ferragina & Manzini, 2000). Taking into account at most *N* − *k* + 1 different *k*-mers, the upper limit is *O*(*N* (*k* + *occ*)) (see Table 9).

**Table 9.** Runtime classes and space occupation module-wise with *N* as the total library size, *n* the number of references per library, *κ*_max_ the largest *k*-mer length, *ω* the window width, and *τ* the amplicon length.

|  | FM Index | FM Frequency | Filter & Transform | Combine |
| --- | --- | --- | --- | --- |
| Runtime | $\mathcal{O}(N)$ | $\mathcal{O}(N(\kappa_{\max} + occ))$ | $\mathcal{O}(\kappa_{\max}N)$ | $\mathcal{O}(n\kappa_{\max}(\frac{N}{n} - \tau_{\min})(\tau_{\max} - \tau_{\min}))$ |
| Space | $\mathcal{O}(N)$ | $\mathcal{O}(N)$ | $\mathcal{O}(N)$ | $\mathcal{O}(n(\frac{N}{n} - \tau_{\min})(\tau_{\max} - \tau_{\min}))$ |

All chemical filters in the **filter & transform** step analyse the encoded sequences in a single- or multi-pass fashion. Some require simple counting (Tm, CG content) or pattern match (dinucleotide runs, (A|T)_3_ tails). The most complex one is the self-annnealing filter in which a *k*-mer is shifted at most 2*k* times and XORed against its complement or reverse complement. The identification of a connected self-annealing pattern of size four (corresponding to a `0b11111111` pattern) can be done in constant time by using bit parallelism (see code on GitHub for details). Filtering is done on at most *N k*-mers, giving us a total runtime of 𝒪(*kN*). For storing the bit vector transformed references, and the ranked TKmerIDs we need *O*(*N*) space. During the **combine** step we additionally store matching *k*-mers. A reference has an expected length of 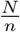 bases. For each forward *k*-mer we search with an offset of *τ*_min_ a candidates’ window of size *τ*_max_ − *τ*_min_. Hence, we have at most 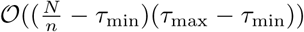 pairs per reference (*κ*_min_, *κ*_max_ are dropped here). For each pair a matchability check is done linear in the expected length of the two *k*-mers, i.e. in 𝒪(*k*) (see *C*_*p*_ in Table 2). Altogether we have *n* references, giving us a total combination runtime of 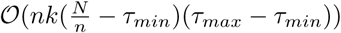.

Space occupancy is linear for the FM index, **frequency**, and **filter & transform** steps. Per text position we use at most one 64 bit unsigned integer independent of how many *k*-mers with varying lengths occur at a specific position (see compression scheme described in Section 3.3.1) and bit vectors for location encoding in the length of the library (see Section 3.3.2). In the **filter & transform** step *k*-mers get cancelled out by resetting length bits in the prefixes of the *k*-mer codes; no additional space is required. When finally combining the remaining set of *k*-mers, we have up to 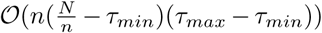 many pairs as described above in detail.

## 5. Discussion

Despite the reference database has a low sequence coverage for plankton taxa, we found some new primer pairs offering a larger coverage or barcode variation than published ones and are chemically suitable for a paired-end PCR (see Table 2). PriSeT correctly output primer pairs that are known to be present in the library if and only if they pass the constraint sets *C*_*s*_ and *C*_*p*_ (see Table 2). When having complete genomes available for primer discovery too many candidates are produced and it is necessary to narrow down the primer sequence constraints or filter in a post-processing step, e.g., for pairs producing amplicons that are distinctive or span exons.

The experiments showed that when searching primer pairs for metabarcoding experiments, it is appropriate to use frequency as an initial filtering heuristic. Only *k*-mers occurring with a minimum frequency will later satisfy sufficient coverage or amplicon variation. The FM index is a transformation that supports frequency queries with lower costs compared to a seed-and-extend approach, e.g. FastPCR by Kalendar et al. (2017), or a MSA-based approach, which requires manageable data sets in order to identify conserved regions serving as primer binding sites.

None of the existing primer search tools that we found is capable of processing multi-sequence libraries and optimizing for frequent primer pairs at the same time. PriSeT is built to fill this gap. Its heuristic approach additionally avoids the necessity to curate an existing library and makes it robust against mislabeled or poor quality references. This in turn gives users more resources to focus on the actual analysis. With sinking costs of NGS, databases are growing on a daily basis, making curation even infeasible and with regard to the sparseness of some clades, we cannot afford to exclude resources.

Since GenBank has no standard specification for labeling sequences by their origin upon upload (e.g. as 18S or COI), our sampling approach also collected non-18S sequences, which explains the relatively low values for coverage and amplicon variation. When a user evaluates PriSeT’s computed primer pairs and their statistics, the heterogeneity of the database has to be taken into consideration.

The PriSeT version at hand does not include coverage or amplicon variation criteria into the filtering for not limiting options – the benefit of a higher coverage is in many cases paid with a lower number of distinct reads (see results in Table 8). We leave it up to the user to decide when coverage is favored over amplicon variation and vice versa. However, the recent coronavirus outbreak demonstrated the importance of scenarios where the goal is to yield barcodes being discriminators of clades.

## 6. Outlook

PriSeT operates batch-wise, i.e. all *k*-mers with frequencies exceeding F are collected at once into a single data structure, filtered chemically, and combined reference-wise. To give an example, Dikarya from the Fungi realm (clade 451864) produces 119.4 million *k*-mers. The main memory occupation of the location map received from GenMap represents the current bottleneck of PriSeT. Processing libraries beyond 500 MB is currently only feasible when increasing the *k*-mer frequency cutoff, s.t. not more than roughly 120 million *k*-mers (≈ 1 Gigabyte) are produced^16^.

In a future PriSeT version this can be tackled by interweaving *k*-mer frequency and filtering: a frequent *k*-mer immediately undergoes filtering, and is only collected when satisfying the frequency threshold F and constraint set *C*_*s*_. This reduces the overall amount of temporarily stored *k*-mers. Input libraries composed of multiple reference sequences would additionally profit from reference-wise partitioning approach. This strategy can be carried on to the **combine** step, since *k*-mers are only combinable if they refer to the same sequence; each of the references is processible in parallel. The current version of PriSeT does not use any thread or process parallelism.

The computationally most expensive part of PriSeT is the **combine** step with a runtime quadratic in *N* where the window size plays in. It is therefore important to set the target read lengths sizes as tight as possible (see Table 9).

Stable binding of primer to template is crucial for the success of a PCR. A single mismatch, especially at the 3’-end, may result in an ineffective PCR. On that account PriSeT is using the (*k*, 0)-frequency to gather only *k*-mer locations with 100 % sequence identity. A future version of PriSeT may allow for up to four errors *e* (or mismatches) for primer sequences in case there are calls for it. When allowing errors, a single *k*-mer occurrence is counted into all location collections associated to *k*-mers with Hamming distances ≤*e*. This has a huge impact on the collection sizes and will have to be chosen carefully.

In metabarcoding we are sometimes interested in primer pairs enclosing barcodes which allow guaranteed distinction of species from clade X from species of another clade Y. If we take as an example the recent SARS-CoV-2 outbreak, one question is, given a sputum sample of a person with flu-like symptoms, does it contain viruses of influenza or corona? We do not want the test to produce false negatives, because the variant is either not sequenced yet or the amplicon has a critical sequence proximity to other viruses. For the *de novo* computation for SARS-CoV-2 submitted online BLAST queries to assure that we access all sequences currently available to GenBank. For a better workflow this feature can be implemented into PriSeT via the NCBI’s Entrez Programming Utilities.

## Supporting information

S1 Primers for Plankton

S2 Primers for SARS-CoV-2

## 7. Additional Files

### 7.1. Supplementary Table — S1_Primers_PriSeT_Plankton.png

Top 1 list of *de novo* sequences per clade computed by PriSeT. All chemical sequence properties are within the constraints listed in Table 2. A more comprehensive list (most 50 frequent primer pairs) can be found on GitHub.

### 7.2. Supplementary File — S2_Primers_PriSeT_SARS-CoV-2.csv

*De novo* primers for SARS-CoV-2 satisfying RT-PCR constraints and producing theoretical amplicons with no occurrences outside the *Sarbecovirus* subspecies. The computation is based on 19 SARS-CoV-2 genomes and the amplicon checks were done online against GenBank’s nr/nt data set.

## Footnotes

1 An exception is the multiplex PCR where multiple non-interfering primer pairs are added.

2 An exception is the multiplex PCR where multiple non-interfering primer pairs are added.

3 www.ncbi.nlm.nih.gov/tools/primer-blast

4 http://bisearch.enzim.hu/?m=genompsearch

5 compared to 1 second by PriSeT (see Figure 6)

6 attempt of accession on 24th February 2020 http://bioinformatics.cribi.unipd.it/primex

7 suffix of length three no displayed

8 https://github.com/mariehoffmann/genmap

9 after applying a frequency cutoff of 5 %

10 Concretely, we use the `sdsl::bit_vector` by Gog et al. (2014) combined with `sdsl::rank_support_v5` and `sdsl::select_support_mcl` occupying at most 0.0625*n*, and 0.2*n* extra bits, respectively (Vigna, 2008; Clark, 1998)

11 accessed on 29.03.2019

12 https://pr2-database.org

13 https://github.com/mariehoffmann/PriSeT_denovo

14 Downloaded from GenBank on 3rd April 2020. Concrete accession identifiers can be requested from the first author.

15 on 3rd of April 2020

16 exemplary for a desktop computer with 16 GB RAM

## References

Benson, D. A., Cavanaugh, M., Clark, K., Karsch-Mizrachi, I., Lipman, D. J., Ostell, J., and Sayers, E. W. GenBank. Nucleic Acids Research, 41(D1):D36–D42, 11 2012. ISSN 0305-1048. doi: 10.1093/nar/gks1195. URL https://doi.org/10.1093/nar/gks1195.

Blaxter, M., Ley, P., Garey, J., Liu, L. X., Scheldeman, P., Vierstraete, A., Vanfleteren, J., Mackey, L., Dorris, M., Frisse, L., Vida, J., and Thomas, W. A molecular evolutionary framework for the phylum nematoda. Nature, 392:71–5, 03 1998. doi: 10.1038/32160.

Boscaro, V., Rossi, A., Vannini, C., Verni, F., Fokin, S. I., and Petroni, G. Strengths and biases of high-throughput sequencing data in the characterization of freshwater ciliate microbiomes. Microbial Ecology, 73:865–875, 2017. doi: 10.1007/s00248-016-0912-8.

Breslauer, K., Frank, R., Blöcker, H., and Marky, L. Predicting dna duplex stability from the base sequence. Proceedings of the National Academy of Sciences of the United States of America, 83:3746–50, 07 1986. doi: 10.1073/pnas.83.11.3746.

Callahan, B., Mcmurdie, P., Rosen, M., Han, A., Johnson, A. J., and Holmes, S. Dada2: High-resolution sample inference from illumina amplicon data. Nature Methods, 13, 05 2016. doi: 10.1038/nmeth.3869.

Cannon, M., Bogale, H., Rutt, L., Humphrys, M., Korpe, P., Duggal, P., Ravel, J., and Serre, D. A high-throughput sequencing assay to comprehensively detect and characterize unicellular eukaryotes and helminths from biological and environmental samples. Microbiome, 6, 12 2018. doi: 10.1186/s40168-018-0581-6.

Clark, D. R. Compact Pat Trees. PhD thesis, CAN, 1998.

Derrien, T., Estellé, J., Marco Sola, S., Knowles, D. G., Raineri, E., Guigó, R., and Ribeca, P. Fast computation and applications of genome mappability. PLOS ONE, 7(1):1–16, 01 2012. doi: 10.1371/journal.pone.0030377. URL https://doi.org/10.1371/journal.pone.0030377.

Elbrecht, V., Braukmann, T. W. A., Ivanova, N. V., Prosser, S. W. J., Hajibabaei, M., Wright, M., Zakharov, E. V., Hebert, P. D. N., and Steinke, D. Validation of coi metabarcoding primers for terrestrial arthropods. PeerJ, 7, 2019. doi: 10.7717/peerj.7745.

Ferragina, P. and Manzini, G. Opportunistic data structures with applications. volume 2000, pp. 390–398, 02 2000. ISBN 0-7695-0850-2. doi: 10.1109/SFCS.2000.892127.

Gog, S., Beller, T., Moffat, A., and Petri, M. From theory to practice: Plug and play with succinct data structures. In 13th International Symposium on Experimental Algorithms, (SEA 2014), pp. 326–337, 2014.

Hadziavdic, K., Lekang, K., Lanzen, A., Jonassen, I., Thompson, E. M., and Troedsson, C. Characterization of the 18s rrna gene for designing universal eukaryote specific primers. PLOS ONE, 9(2):1–10, 02 2014. doi: 10.1371/journal.pone.0087624. URL https://doi.org/10.1371/journal.pone.0087624.

Harder, C., Rønn, R., Brejnrod, A., Bass, D., Abu Al-Soud, W., and Ekelund, F. Local diversity of heathland cercozoa explored by in-depth sequencing. The ISME Journal, 10, 03 2016. doi: 10.1038/ismej.2016.31.

Kalendar, R., Khassenov, B., Ramankulov, Y., Samuilova, O., and Ivanov, K. I. Fastpcr: An in silico tool for fast primer and probe design and advanced sequence analysis. Genomics, 109(3):312 – 319, 2017. ISSN 0888-7543. doi: https://doi.org/10.1016/j.ygeno.2017.05.005. URL http://www.sciencedirect.com/science/article/pii/S0888754317300368.

Kämpke, T., Kieninger, M., and Mecklenburg, M. Efficient primer design algorithms. Bioinformatics (Oxford, England), 17:214–25, 04 2001. doi: 10.1093/bioinformatics/17.3.214.

Lexa, M. and Valle, G. Primex: Rapid identification of oligonucleotide matches in whole genomes. Bioinformatics (Oxford, England), 19:2486–8, 01 2004. doi: 10.1093/bioinformatics/btg350.

Medlin, L., Elwood, H. J., Stickel, S., and Sogin, M. L. The characterization of enzymatically amplified eukaryotic 16s-like rrna-coding regions. Gene, 71(2):491 – 499, 1988. ISSN 0378-1119. doi: https://doi.org/10.1016/0378-1119(88)90066-2. URL http://www.sciencedirect.com/science/article/pii/0378111988900662.

Moreno, Y., Moreno-Mesonero, L., Amorós, I., Pérez, R., Morillo, J., and Alonso, J. Multiple identification of most important waterborne protozoa in surface water used for irrigation purposes by 18s rrna amplicon-based metagenomics. International Journal of Hygiene and Environmental Health, 221(1):102 – 111, 2018. ISSN 1438-4639. doi: https://doi.org/10.1016/j.ijheh.2017.10.008. URL http://www.sciencedirect.com/science/article/pii/S1438463917305850.

Pockrandt, C., Alzamel, M., Iliopoulos, C. S., and Reinert, K. Genmap: Fast and exact computation of genome mappability. bioRxiv, 2019. doi: 10.1101/611160.

Rappé, M. S. and Giovannoni, S. J. The uncultured microbial majority. Annual Review of Microbiology, 57(1): 369–394, 2003. doi: 10.1146/annurev.micro.57.030502. 090759. URL https://doi.org/10.1146/annurev.micro.57.030502.090759. PMID: 14527284.

Schmidt, T., Rodrigues, J., and von Mering, C. Ecological consistency of ssu rrna-based operational taxonomic units at a global scale. PLoS computational biology, 10: e1003594, 04 2014. doi: 10.1371/journal.pcbi.1003594.

Smart, A. S., Tingley, R., Weeks, A. R., van Rooyen, A. R., and McCarthy, M. A. Environmental dna sampling is more sensitive than a traditional survey technique for detecting an aquatic invader. Ecological Applications, 25(7):1944–1952, 2015. doi: 10.1890/14-1751.1. URL https://esajournals.onlinelibrary.wiley.com/doi/abs/10.1890/14-1751.1.

Smith, K. F., Kohli, G. S., Murray, S. A., and Rhodes, L. L. Assessment of the metabarcoding approach for community analysis of benthic-epiphytic dinoflagellates using mock communities. New Zealand Journal of Marine and Freshwater Research, 51(4):555–576, 2017. doi: 10.1080/00288330.2017.1298632. URL https://doi.org/10.1080/00288330.2017.1298632.

Stoeck, T., Bass, D., Nebel, M., Christen, R., Jones, M. D. M., Breiner, H.-W., and Richards, T. A. Multiple marker parallel tag environmental dna sequencing reveals a highly complex eukaryotic community in marine anoxic water. Molecular Ecology, 19(1):21–31, 2010. doi: 10.1111/j.1365-294X.2009.04480.x. URL https://onlinelibrary.wiley.com/doi/abs/10.1111/j.1365-294X.2009.04480.x.

Tusnády, G. E., Simon, I., Váradi, A., and Arányi, T. BiSearch: primer-design and search tool for PCR on bisulfite-treated genomes. Nucleic Acids Research, 33 (1):e9–e9, 01 2005. ISSN 0305-1048. doi: 10.1093/nar/gni012. URL https://doi.org/10.1093/nar/gni012.

Valiente Moro, C., Crouzet, O., Rasconi, S., Thouvenot, A., Coffe, G., Batisson, I., and Bohatier, J. New design strategy for development of specific primer sets for pcr-based detection of chlorophyceae and bacillariophyceae in environmental samples. Applied and Environmental Microbiology, 75(17):5729–5733, 2009. ISSN 0099-2240. doi: 10.1128/AEM.00509-09. URL https://aem.asm.org/content/75/17/5729.

Vigna, S. Broadword implementation of rank/select queries. In McGeoch, C. C. (ed.), Experimental Algorithms, pp. 154–168, Berlin, Heidelberg, 2008. Springer Berlin Heidelberg. ISBN 978-3-540-68552-4.

Visco, J. A., Apothéloz-Perret-Gentil, L., Cordonier, A., Esling, P., Pillet, L., and Pawlowski, J. Environmental monitoring: Inferring the diatom index from next-generation sequencing data. Environmental Science & Technology, 49(13):7597–7605, 2015. doi: 10.1021/es506158m. URL https://doi.org/10.1021/es506158m. PMID: 26052741.

Ye, J., Coulouris, G., Zaretskaya, I., Cutcutache, I., Rozen, S., and Madden, T. Primer-blast: a tool to design target-specific primers for polymerase chain reaction. bmc bioinform 13:134. BMC bioinformatics, 13:134, 06 2012. doi: 10.1186/1471-2105-13-134.

Yoon, T.-H., Kang, H.-E., Kang, C.-K., Lee, S. H., Ahn, D.-H., Park, H., and Hyun-Woo, K. Development of a cost-effective metabarcoding strategy for analysis of the marine phytoplankton community. PeerJ, 4:e2115, 2016. doi: 10.7717/peerj.2115.

Zhou, P., Yang, X., Wang, X.-G., Hu, B., Zhang, L., Zhang, W., Si, H.-R., Zhu, Y., Li, B., Huang, C.-L., Chen, H.-D., Chen, J., Luo, Y., Guo, H., Jiang, R.-D., Liu, M.-Q., Chen, Y., Shen, X.-R., and Wang, X. A pneumonia outbreak associated with a new coronavirus of probable bat origin. Nature, 579, 03 2020. doi: 10.1038/s41586-020-2012-7.

Zimmermann, J., Jahn, R., and Gemeinholzer, B. Barcoding diatoms: Evaluation of the v4 subregion on the 18s rrna gene, including new primers and protocols. Organisms Diversity & Evolution, 11:173–192, 07 2011. doi: 10.1007/s13127-011-0050-6.

