## Supplementary material for "PriSeT: Efficient *De Novo* Primer Discovery": S1 Primers for Plankton

| Primer | Sequence (fwd/rev) | Tm | CG |
|---|---|---|---|
| 1426f6f9b97f501a | 3-AAACGGCTACCACATCC-> | 52 | 0.53 |
| | <-GGTATTTGCTACGGCTG-3 | 52 | 0.53 |
| 24555f6837b9651f | 3-TTCTTAGTTGGTGGAGTG-> | 52 | 0.44 |
| | <-GTCCAGACACTACGGGA-3 | 54 | 0.59 |
| 2afb9ade5ef548f | 3-ACAGGGAGGTAGTGACA-> | 52 | 0.53 |
| | <-CTTGCTTTCAATCCCCTA-3 | 52 | 0.44 |
| 33c14baf2ac76276 | 3-GGCTACCACATCCAAGG-> | 54 | 0.59 |
| | <-CTTGCTTTCAATCCCCTA-3 | 52 | 0.44 |
| 35050c2634d666c7 | 3-GGCTACCACATCCAAGG-> | 54 | 0.588 |
| | <-GTGTCCCTCCATCACTGTT-3 | 58 | 0.526 |
| 38f9227a340f05e | 3-GCCGTTCTTAGTTGGTG-> | 52 | 0.53 |
| | <-GTCCAGACACTACGGGA-3 | 54 | 0.59 |
| 3f32736b9d094915 | 3-TGAAACACGGACCAAGG-> | 52 | 0.53 |
| | <-CTCCTTTGAGACCACCT-3 | 52 | 0.53 |
| 412d47502c9c1b | 3-TTCTTAGTTGGTGGAGTG-> | 52 | 0.44 |
| | <-TCTACAAGACCCGGCGT-3 | 54 | 0.59 |
| 4dbdfed2605e324 | 3-CCCTGTCCTCCTATTTTC-> | 54 | 0.5 |
| | <-CTACGGCTGGTCGCTAA-3 | 54 | 0.59 |
| 57bc43fe1080644d | 3-TAGGTGCCGAGAAGATG-> | 52 | 0.53 |
| | <-CACTTGGACGCCTTCCT-3 | 54 | 0.59 |
| 5f5fdc94bf3f7db4 | 3-TTCTTAGTTGGTGGAGTG-> | 52 | 0.44 |
| | <-CCTTCAAACTCCGTTATTG-3 | 54 | 0.42 |
| 6f74c4dbf4ceae12 | 3-TCTGCCAAGGATGTTTTC-> | 52 | 0.44 |
| | <-CTACGGCTGGTCGCTAA-3 | 54 | 0.59 |
| 7aab6403d6856205 | 3-AAATCTCAACGCATACTGC-> | 54 | 0.42 |
| | <-CGAAACATAGGGCGAAC-3 | 52 | 0.53 |
| 7e079b504409f8c0 | 3-ACCGTGAGGGAAAGATG-> | 52 | 0.53 |
| | <-ACTTTGTGCCTGGTTCC-3 | 52 | 0.53 |
| 8d14274d691d15e5 | 3-GGCTACCACATCCAAGG-> | 54 | 0.59 |
| | <-CATACCAGCGTTCCGAC-3 | 54 | 0.59 |
| 96c04483d9557b08 | 3-CACAGGGAGGTAGTGAC-> | 54 | 0.59 |
| | <-ACCTCCCGTTCAGACCA-3 | 54 | 0.59 |
| af8f7968a5ee777e | 3-GGTCTGTGATGCTCCTC-> | 54 | 0.59 |
| | <-CCTTACAGAGCATCCGC-3 | 54 | 0.59 |
| b099967d0f5ac180 | 3-GTGAGGATTGACAGATTG-> | 52 | 0.44 |
| | <-GGCAAGAATCAACCACC-3 | 52 | 0.53 |
| b7488789aaf96d7b | 3-TGGTGGGTTGCCTTGTC-> | 54 | 0.59 |
| | <-GTCCAGACACTACGGGA-3 | 54 | 0.59 |
| b805fd5bf8167cc7 | 3-GATACCTTCCTCAATCAAG-> | 54 | 0.42 |
| | <-ATCTCTGGTAACATCAGG-3 | 52 | 0.44 |
| bbcb9dc15a5fc34c | 3-GTGAACCTGCGGAAGGAT-> | 56 | 0.56 |
| | <-TGTGTTGGGATACGCCA-3 | 52 | 0.53 |
| c2ee8143ec040d5a | 3-AGGGAGGTAGTGACAAG-> | 52 | 0.53 |
| | <-AACTGCCTTCCCGTGGT-3 | 54 | 0.59 |
| cb6c82029453de6a | 3-GGCTACCACATCCAAGG-> | 54 | 0.59 |
| | <-TGGCAGCATCAGAGTTG-3 | 52 | 0.53 |
| cce22a01d74086cd | 3-GGCTACCACATCCAAGG-> | 54 | 0.59 |
| | <-CCTCATACCAGCGTTCC-3 | 54 | 0.59 |
| d8d47dc9b873d02b | 3-GGCTACCACATCCAAGG-> | 54 | 0.59 |
| | <-CTTTCGTAAACGGTTCCT-3 | 52 | 0.44 |
| d90e133cfe247ad5 | 3-GGCTACCACATCTAAGG-> | 52 | 0.53 |
| | <-AACTGCCTTCCCGTGGT-3 | 54 | 0.59 |
| eb59a790ece1766 | 3-AGCACCTTGTGAGAAATC-> | 52 | 0.44 |
| | <-AACTAAGATACCCACCAC-3 | 52 | 0.44 |
| ecb3cb9fe242b24e | 3-GTGTCGCTCTTCTTCCC-> | 54 | 0.59 |
| | <-CGAAACATAGGGCGAAC-3 | 52 | 0.53 |
| edcd39cccec9d72a | 3-AACACGGACCAAGGAGT-> | 52 | 0.53 |
| | <-GGCTTTCTACCACTTGAT-3 | 52 | 0.44 |
| f83f6dae50175268 | 3-TGCGGCTTAATTTGACTC-> | 52 | 0.44 |
| | <-CACGTACCGGCAAGAATC-3 | 56 | 0.56 |
| fcb8901444edb4b0 | 3-ACACGGGGAAACTTACC-> | 52 | 0.53 |
| | <-GGCAAGAATCAACCACCT-3 | 54 | 0.5 |
